# Compact refractive dual-channel AOSLO for wide-field imaging in mice reveals microglial interactions with transplanted neurons

**DOI:** 10.1101/2025.03.31.645335

**Authors:** Zhongqiang Li, Stella Mary, Thomas V Johnson, Ji Yi

## Abstract

Adaptive optics (AO) enables cellular-resolution retinal imaging, yet mirror-based AOSLO systems are constrained by off-axis aberrations that restrict high-quality imaging to narrow fields of view, limiting in vivo studies of dynamic, large-scale retinal processes. Clinical translation of regenerative cell therapy to neurologic disease is hampered by attrition of donor neurons following transplantation. We hypothesized that early innate neuroinflammatory responses to retinal ganglion cell (RGC) transplantation underlie early death of donor cells and that next-generation imaging technologies would provide evidence for microglial attack of grafted neurons. We present a compact refractive lens-based AOSLO system that achieves two-color fluorescence imaging across up to a 16° field of view in mouse retina. Dual-wavelength excitation (488 nm and 552 nm) enables visualization of two fluorescence labels, while AO correction improves axial resolution and depth fidelity, allowing robust separation of structures through anatomical layers in retina. Using this platform, we performed 3D time-lapse imaging of microglia and longitudinal imaging in an optic nerve crush model, revealing layer-dependent differences in microglial motility, early activation signatures, and large-scale redistribution longitudinally. The system enabled widefield visualization of injury-associated vascular changes and spatial coupling between microglia and vasculature. Finally, depth-resolved two-color imaging captured immune responses to intravitreally transplanted RGCs, including host-cell recruitment, rapid neurite retraction following local immune-cell contact, and microglial phagocytosis of donor RGCs. Together, these results demonstrate that refractive AOSLO enables in vivo observations of microvascular organization, neuroimmune dynamics, injury responses, and transplanted-cell behavior with spatiotemporal resolution. Our data also suggests that modulation of microglial reactivity may improve outcomes of RGC transplantation.

**Significance Statement:** Wide-field, depth-resolved imaging is essential for understanding how different cell types interact across a large retinal area, yet existing AOSLO systems retain limited imaging fields due to a conventional optical design using reflective spherical mirrors. Our refractive large-FOV AOSLO overcomes this limitation, enabling simultaneous two-color, diffraction-limited 3D imaging across an up to 16° field in vivo. This platform reveals previously inaccessible biological phenomena—including layer-specific microglial dynamics after optic nerve injury, microvascular remodeling, and rapid microglial rejection of transplanted RGCs—providing critical insight into neuroimmune behavior and retinal repair mechanisms at single-cell and subcellular resolution.

## Introduction

The retina is directly accessible using light, providing a unique window into neural structure, cellular interactions, and disease progression *in vivo*. Beyond its central role in vision, the retina serves as a powerful model system for studying neurodegeneration, neuroinflammation, and emerging cell-based regenerative therapies (1). Many retinal and optic neuropathies—including glaucoma and traumatic optic nerve injury—are characterized by retinal ganglion cell (RGC) degeneration accompanied by profound neuroinflammatory responses, particularly involving microglia (2–5). Understanding how neuronal injury, neuroimmune activation, and neuronal replacement strategies unfold over time requires imaging tools capable of resolving cellular and subcellular dynamics across large spatial areas and multiple retinal layers.

Advances in retinal imaging have substantially improved our understanding of retinal anatomy and pathology (6, 7). Conventional imaging modalities, such as fundus photography, optical coherence tomography (OCT), and scanning laser ophthalmoscopy (SLO), provide important structural and functional information but remain fundamentally limited in spatial resolution by the eye’s intrinsic optical aberrations (1, 6–8). These limitations are particularly restrictive for studying neuroimmune interactions, such as morphological microglial remodeling, cell migration, and neurite dynamics, which occur at micrometer scales and evolve dynamically over time. Similarly, investigations of optic neuropathy pathogenesis and retinal ganglion cell transplantation demand imaging approaches that can capture both early localized responses and later large-scale tissue remodeling.

Adaptive optics (AO) imaging corrects ocular wavefront aberrations and enables near–diffraction-limited retinal imaging *in vivo*, allowing visualization of individual retinal cells and subcellular structures (8–11). AO scanning laser ophthalmoscopy (AOSLO), originally adapted from astronomical applications, has enabled detailed imaging of photoreceptors, RGCs, microvasculature, and immune cells in animal models and humans (6, 7, 9, 12–14). However, most existing AOSLO systems are optimized for small diffraction-limited fields of view (around 1° to 3°(15)) and predominantly two-dimensional imaging, limiting their ability to capture spatially distributed cellular responses, depth-dependent interactions, and time-evolving behaviors (15–17). These constraints pose a major challenge for studying biological processes such as microglial activation after injury, layer-specific immune redistribution, and the migration and integration of transplanted neurons, all of which span hundreds of micrometers in depth and millimeters in lateral extent.

Refractive lens-based AOSLO and AOSLO/OCT systems have emerged as an alternative technology, offering compact designs, including handheld implementations, and a slightly larger diffraction-limited FOV (as large as 4° to 5°) (15, 18–27). However, refractive systems face challenges arising from lens surface reflections that interfere with wavefront sensing and chromatic aberrations that render wavefront distortion wavelength dependent (28). Both computational approaches, such as sensorless AO, and optical strategies, including polarization-based rejection of surface reflections, have been proposed to address these limitations (20). Importantly, for studies requiring dynamic, multicolor fluorescence imaging— such as simultaneously tracking resident immune cells and transplanted neurons—robust correction across wavelengths and depth is essential.

Here, we present a compact, lens-based AOSLO system designed to address these unmet biological imaging needs. The system enables simultaneous, volumetric two-color fluorescence imaging (e.g., 488 nm and 552 nm) in the mouse retina while maintaining near–diffraction-limited performance. By incorporating polarization optics within a conventional Shack–Hartmann (SH) wavefront sensing scheme and using excitation lasers for channel-specific AO correction, the system effectively suppresses lens surface reflections and mitigates chromatic aberration effects. Importantly, the system supports imaging over a field of view of up to 20°. In the experiments reported here, a 16° field of view was used to ensure stable aberration correction and consistent image quality across the field. This configuration enables wide-area surveys to identify regions of interest, followed by high-resolution, depth-resolved imaging of cellular and subcellular dynamics.

Using this platform, we perform *in vivo* fluorescence angiography, dynamic microglial imaging in healthy and optic nerve–injured retinas, and longitudinal observation of transplanted RGCs and their interactions with host neuroimmune cells. By combining large FOV coverage, true three-dimensional imaging, and time-resolved observation, this system enables direct visualization of how microglial morphology evolves after injury, how immune responses redistribute across retinal layers, and how transplanted neurons migrate and stabilize within the host tissue. Together, these capabilities establish a biologically driven imaging framework for studying retinal neuroinflammation, injury response, and cell-based regenerative strategies *in vivo*.

## Results

### DualCH-AOSLO enables large-FOV retinal angiography with local AO correction

To quantitatively evaluate the performance of adaptive optics (AO) correction, we used a 3-mm N-BK7 ball lens to emulate the mouse eye, providing comparable size and numerical aperture (Supplementary Fig. S1). In the red channel, the lateral and axial resolutions without AO (non-AO) were measured as 1.2 μm and 8.75 μm, respectively (Supplementary Fig. S1A). With AO correction, these improved to 0.8 μm laterally and 4.62 μm axially, representing a >2-fold enhancement in axial resolution and approaching the theoretical diffraction limits of 0.73 μm (lateral) and 4.91 μm (axial). Similarly, in the green channel, AO correction improved the lateral and axial resolutions from 1.56 μm and 10.95 μm (non-AO) to 1.21 μm and 5.40 μm, respectively (Supplementary Fig. S1B, D), compared with theoretical limits of 0.64 μm and 4.29 μm. Notably, the green channel exhibited slightly reduced performance compared to the red channel, suggesting residual chromatic aberration, likely due to system alignment being optimized for the red channel.

The DualCH-AOSLO system (Fig. 1A) enables acquisition of a single raster scan covering up to ~1 × 1 mm^2^ of the mouse retina, corresponding to a maximum achievable field of view (FOV) of ~20°. This wide-field capability enables rapid retinal surveys for identifying regions of interest. Fluorescence angiography was performed using retroorbital injection of fluorescein dextran (fluorescein angiography, FA) or rhodamine dextran (rhodamine angiography, RA) (18). By stitching multiple scans, retinal regions up to ~2.5 × 2.5 mm^2^ were visualized (Fig. 1B-C), enabling large-area mapping of the retinal vasculature and facilitating navigation to smaller regions for subsequent high-resolution AO imaging.

**Figure 1.**
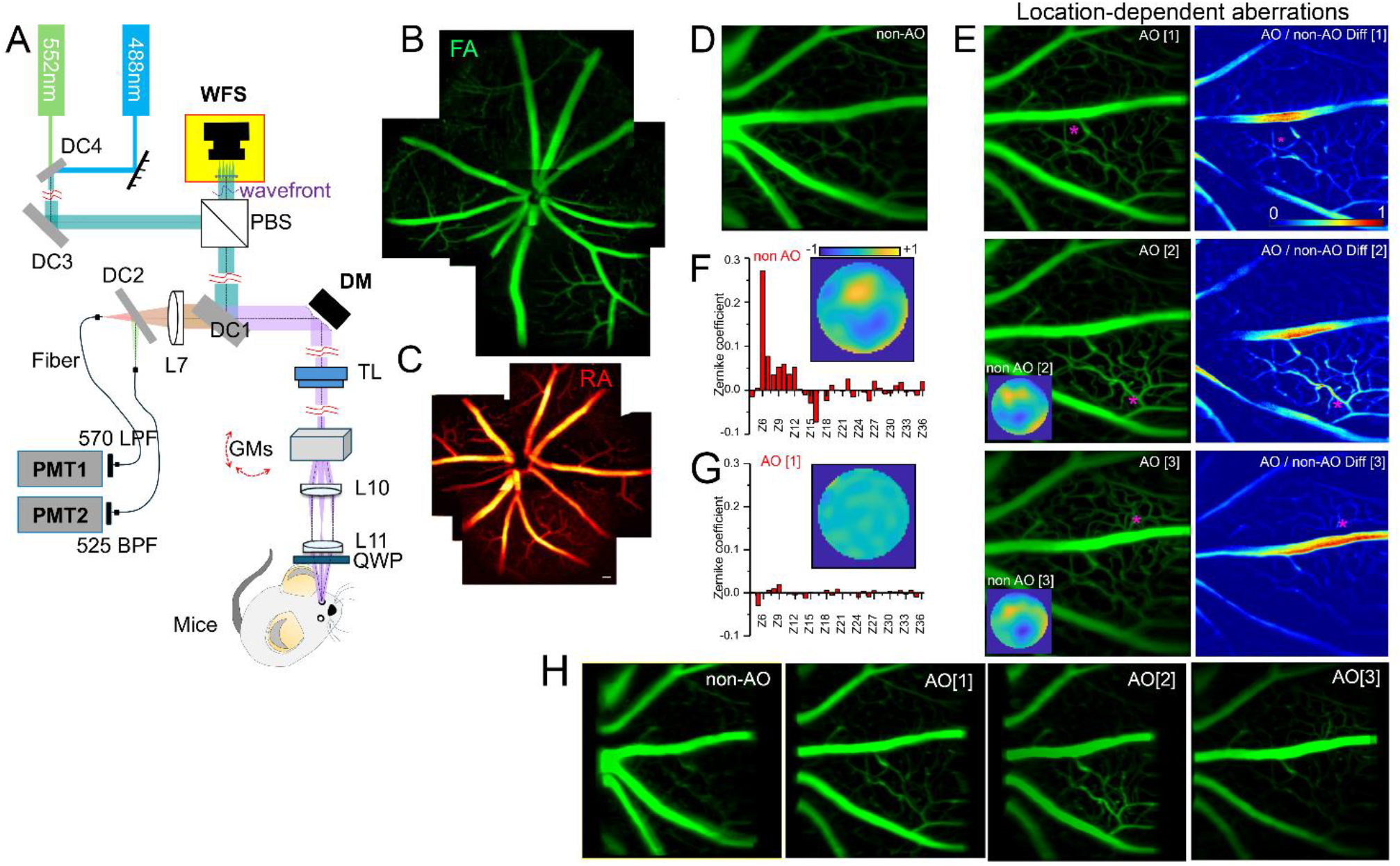
System schematic diagram of DualCH-AOSLO and the Rhodamine-B angiography (RA) and Fluorescein angiography (FA) of DualCH-AOSLO with and without AO correction and their evaluation under large field and small field wavefront sensing. (A) System schematic diagram of DualCH-AOSLO. (B) and (C) The extra-large SLO imaging with the conventional angiography. (D) and (E) The MIPs of FA image stacks (1100×1100×100 μm^3^) without AO correction and with AO corrections performed at different locations. Red asterisk: center of 200×200 μm^2^ WFS area (NO-AO [2] and NO-AO [3] WF maps were inserted in their AO images). (F) and (G) The normalized pixel intensity of AO and NO AO correction under large and small imaging fields in FA. (H) Representative volumetric views of the same retinal region under non-AO and AO conditions: AO[1]-[3]. Compared with non-AO imaging, AO-corrected data reveal improved structural continuity and finer microvascular detail. These views provide a qualitative three-dimensional perspective of the imaged volume. Scale bar: 100 μm.

To evaluate field-dependent aberration correction across the wide imaging field, AO was applied at different positions within the superficial vascular plexus. Without AO, capillary structures were poorly resolved across the FOV (Fig. 1D). When AO correction was performed near the center of the FOV (AO[1]), capillary features were clearly resolved. When the wavefront sensing region was shifted toward the bottom-right (AO[2]) or top-left (AO[3]) of the FOV, localized improvements in signal and capillary clarity were observed at the sensing regions (Fig. 1E). Intensity difference maps (AO/non-AO) revealed spatially localized signal enhancement corresponding to the wavefront measurement locations. Zernike wavefront maps further confirmed effective aberration correction, resulting in flatter residual wavefronts following AO correction (Fig. 1F-G) WFS performance was further evaluated using different sensing field sizes under both FA and RA conditions. Three-dimensional volumetric views further illustrate the improved structural definition with AO correction (Fig. 1H, with rotating renderings provided in Supplementary Videos 1–4).

With a large WFS area (0.8 × 0.8 mm^2^, corresponding to ~16° FOV), AO imaging resolved capillary structures more clearly than non-AO imaging for both contrast agents (Supp. Fig. S2A). When the WFS area was reduced to ~200 × 200 μm^2^, individual capillary morphology appeared noticeably sharper under AO correction in both RA and FA images (Supp. Fig. S2A–D). This improvement was further supported by normalized pixel intensity profiles, which showed higher peak values in AO images than in the corresponding non-AO images (Supp. Fig. S2E–H). Quantitatively, AO correction produced an approximately two-fold increase in normalized signal intensity relative to non-AO imaging in both the large- and small-FOV WFS configurations (Supp. Fig. S2E–H), demonstrating a robust enhancement in vascular contrast across imaging conditions.

We further performed *in vivo* volumetric (3D) analysis of RA and FA imaging to evaluate AO-enabled vascular layer discrimination. In both modalities, AO correction enabled clearer visualization of the three retinal vascular plexuses in *en-face* and cross-sectional views (Supp. Fig. S3A,D).

This improvement was further supported by quantitative intensity profile analysis. Notably, when the analyzed region was positioned within the IVP, the adjacent capillary-like structure originated from the DVP appeared in the IVP image because of axial signal overlap under non-AO imaging (Supp. Fig. S3B,E). In the lateral direction, non-AO imaging showed substantial overlap from the DVP signal, with the normalized intensity remaining high (>0.75–1.0) in regions where this overlapping structure was present. In contrast, under AO correction, the DVP-derived overlapping structure was largely eliminated from the IVP image, and the corresponding normalized intensity dropped to <0.25, indicating markedly improved separation between IVP and DVP signals (Supp. Fig. S3B,E).

In the axial direction, AO imaging resolved the SVP and IVP as two distinct peaks in the normalized intensity profiles, demonstrating successful separation of these two vascular layers (Supp. Fig. S3C,F). By comparison, the corresponding non-AO profiles merged into a single broadened peak, indicating axial overlap and insufficient layer discrimination. Comparable improvements were observed in both RA and FA imaging, confirming that AO substantially enhances depth-resolved vascular separation across imaging modalities.

### Volumetric in vivo imaging resolves layer-specific microglial morphology and 3D time-lapse imaging reveals layer-dependent microglial dynamics

Microglia are critical regulators of CNS equilibrium and response to disease, and their morphology is closely linked to their activation state and physiology. Though characteristics of morphology and dynamics have been studied extensively, work to date has been limited to static time points, *ex vivo* imaging paradigms, or time-lapse *in vivo* imaging that fails to parse microglia according to retinal layer, due to insufficient axial resolution (29–33). To visualize the global distribution of retinal microglia, we first performed wide-field fluorescence fundus imaging using the Micron V imaging system in Cx3cr1-GFP mice. As shown in Fig. 2A, GFP-labeled microglia were broadly distributed across the retina and appeared as scattered punctate signals throughout the field of view. To better understand retinal layer-specific microglial responses under normal and stressed conditions, we leveraged simultaneous DualCH-AOSLO to image *Cx3xr1-GFP* mice using RA as a depth reference (Fig. 2B). Notably, the Micron and AOSLO images shown in panels A and B were acquired from different eyes, illustrating representative imaging results obtained with the two modalities. Microglia were visualized across multiple retinal layers, including the nerve fiber layer (NFL), inner plexiform layer (IPL), and outer plexiform layer (OPL), corresponding to the SVP, IVP, and DVP, respectively (Fig. 2C).

**Figure 2.**
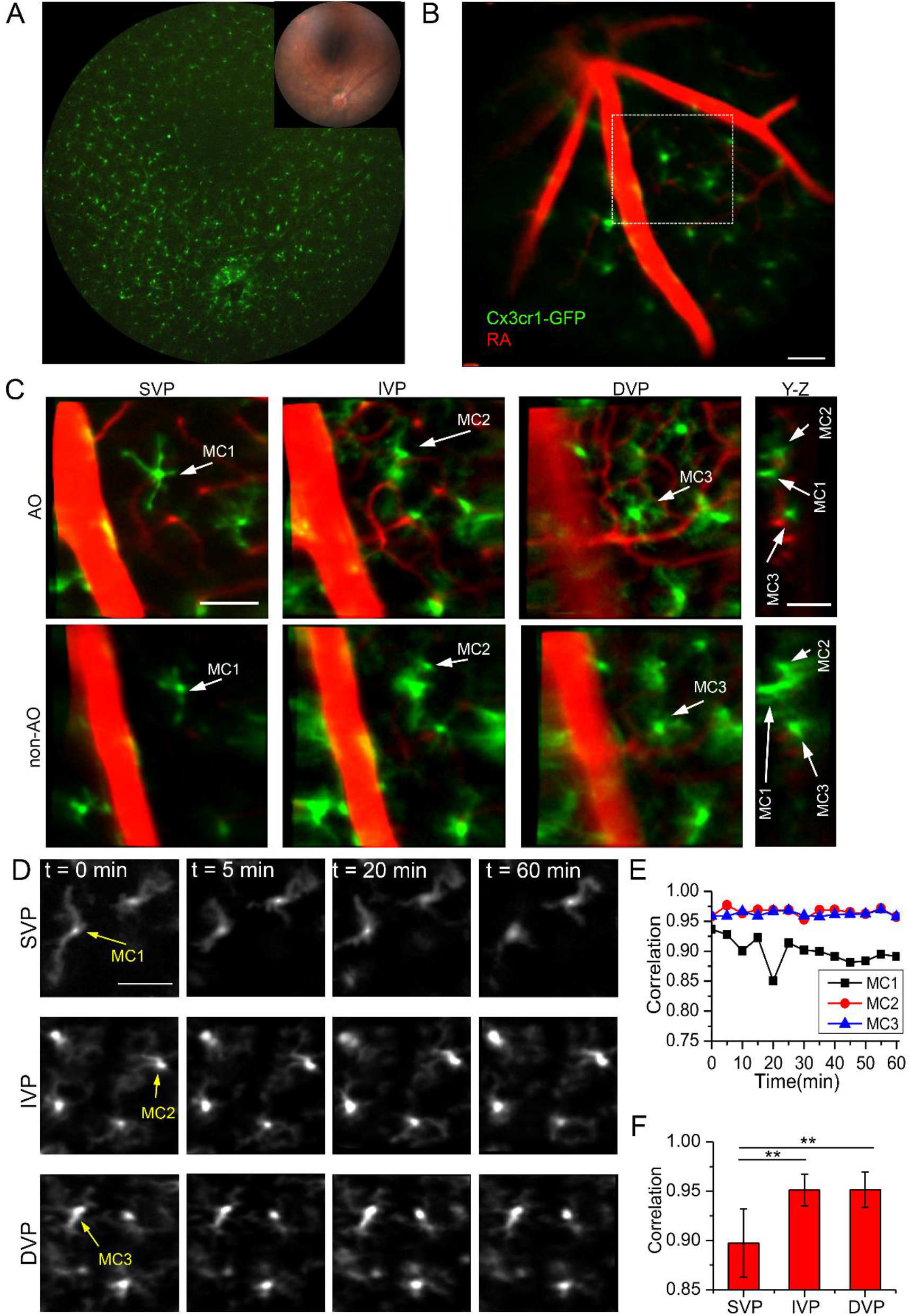
*In-vivo* 3D imaging of microglia cells composited with angiography and their processes dynamics under the AO and NO AO imaging. (A) Wide-field Micron fluorescence fundus image showing the distribution of Cx3cr1-GFP microglia across the retina. (B) Large field conventional SLO images of MCs composited with RA imaging. Images in panels A and B were acquired from different eyes. (C) Three MCs layers distribute along with the three vascular plexus layers under AO imaging and NO AO imaging. (D) MCs with different processes movements at the SVP, IVP and DVP layers. (E) Correlation of individual MC from three different layers at different time points. (F) Correlation of multiple MCs from three different layers among different mice (N= 6 different right eyes, and n= 32 MCs in total). Scale bar: 100 μm.

With AO correction, microglial processes appeared sharper across all layers, with fine branches clearly resolved (Fig. 2C). In contrast, non-AO imaging produced blurred microglial processes, and small branches were not distinguishable (Fig. 2C). Under non-AO conditions, the IVP and DVP layers were poorly separated, whereas AO correction enabled clear axial discrimination between these layers (Fig. 2C). Quantitative intensity profile analysis further confirmed the improvement provided by AO correction (Supp. Fig. S4A). In both the SVP and DVP, microglial processes imaged with AO exhibited consistently narrower lateral and axial intensity profiles than those acquired without AO, indicating improved confinement of the signal in both transverse and axial dimensions. Consistent with these profiles, the lateral resolution was improved by >2x in lateral dimension, and up to 3-4x in axial dimension (Supp. Fig. S4B). These results demonstrate that AO correction markedly enhances both transverse and axial resolution, enabling improved visualization of fine microglial morphology.

Volumetric 3D AOSLO imaging enables time-lapse monitoring of microglial dynamics *in vivo*. In a healthy eye imaged at 5-minute intervals over 60 minutes, microglia in the SVP exhibited dynamic process extension and retraction (Fig. 2D; Supp Fig. S4C). Quantitative analysis of those processes from MC1 showed an average process retraction velocity of 3.43 ± 0.19 μm/min. Over the imaging period, major processes retracted, new branches emerged, and some branches disappeared.

In contrast, microglia located in the deeper IPL and OPL layers displayed minimal morphological changes during the same imaging period (Fig. 2D, Supplementary Videos 5-7; Supp Fig. S4C). Correlation analysis of individual microglial images over time showed lower correlation values for SVP microglia compared with those in the IVP and DVP (Fig. 2E). Across 32 cells, the average correlation for SVP microglia was 0.897 ± 0.035, which was significantly lower than that of IVP microglia (0.951 ± 0.016, *p = 0*.*0028*) and DVP microglia (0.951 ± 0.016, *p = 0*.*0027*) (Fig. 2F).

### Longitudinal in vivo imaging captures microglial responses following optic nerve crush

To understand how layer-specific microglial dynamics differ in the neurodegenerative disease state, DualCH-AOSLO was applied to an optic nerve crush (ONC) mouse model. During the first three days after ONC, microglial morphology in the SVP showed minimal change, while microglial density increased (Fig. 3A; Supp Fig. S5). By day 8 post-ONC, microglia exhibited radial morphology with reduced process complexity in the SVP. These features persisted through day 21, although overall microglial density declined over that timeframe (Fig. 3A).

**Figure 3.**
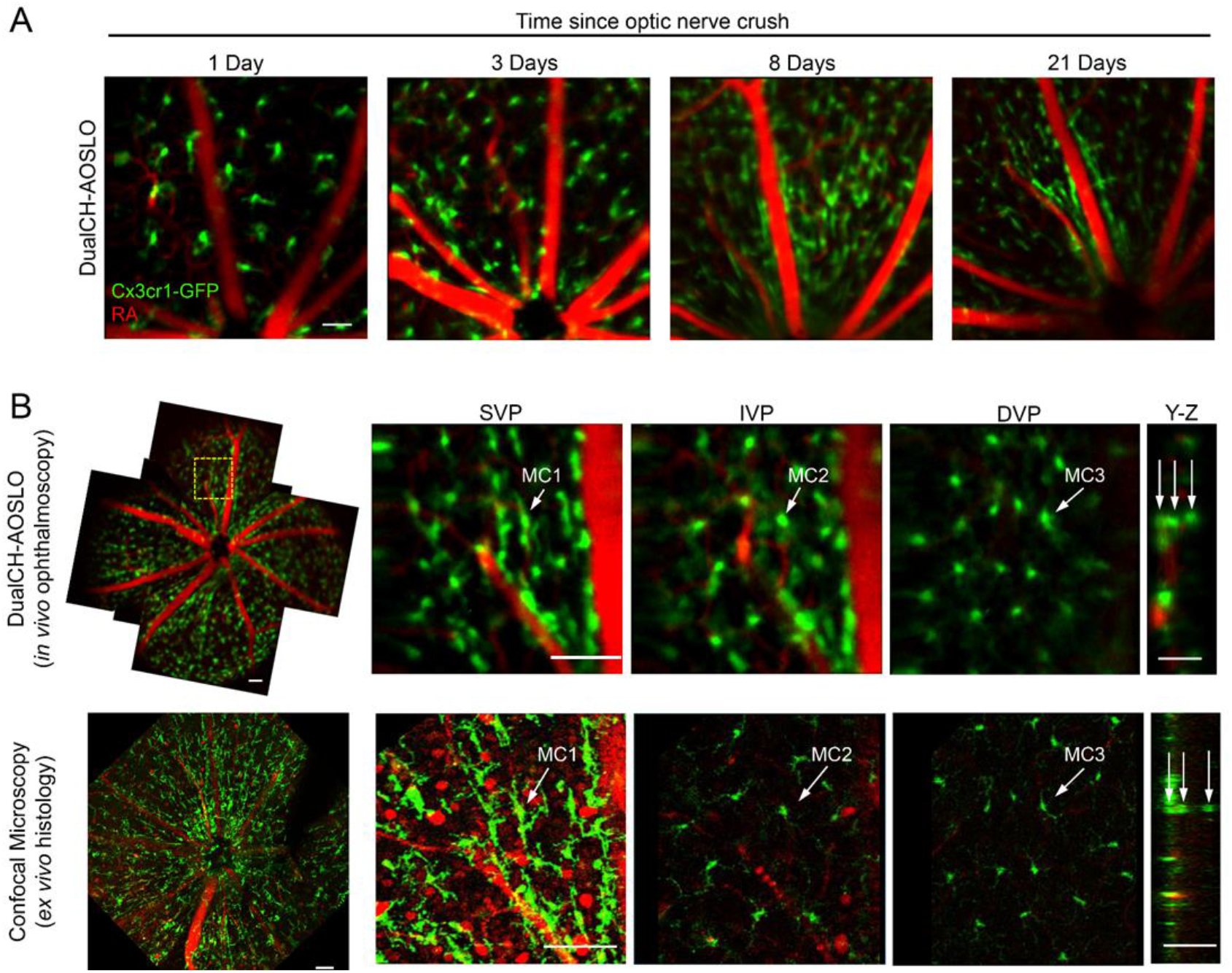
MCs change after optics nerve crush. (A) The *in-vivo* imaging of MCs morphology at SVP on day 1, day 3, day 8, and day 21 after ONC. (B) Wide field of conventional SLO images and its 3D imaging MC morphology on day 21 and compared with *ex-vivo* confocal imaging. Scale bar: 100 μm.

Microglia in the IVP and DVP displayed increased process complexity at later stages (Supp Fig. S5). Radially oriented microglia in the SVP were most prominent near the optic disc and decreased with distance from the disc (Fig. 3B). *Ex vivo* confocal imaging at day 21 confirmed the *in vivo* observations, including radial microglial morphology and layer-specific distribution (Fig. 3B). Histological analysis further showed reduced RGC density in ONC eyes compared with controls (Supp Fig. S6).

### 3D in-vivo imaging reveals microglial recruitment and stage-dependent interactions with transplanted RGCs

RGC transplantation holds potential for restoring vision in optic neuropathies, provided donor neurons survive long term and engraft into the recipient retina. Prior research has demonstrated that the majority of donor RGCs die shortly after transplantation, and microglia are implicated in this attrition (2, 34, 35). Therefore, we utilized depth-resolved DualCH-AOSLO to image neuroimmune responses to transplanted RGCs across different eyes at defined post-transplantation timepoints and imaging locations (Fig. 4A). We transplanted 4×10^5^ human stem cell derived RGCs into the eyes of Cx3cr1-GFP mice two weeks following ILM digestion with intravitreal pronase-E, which we have previously shown is necessary for engraftment (36).

**Figure 4.**
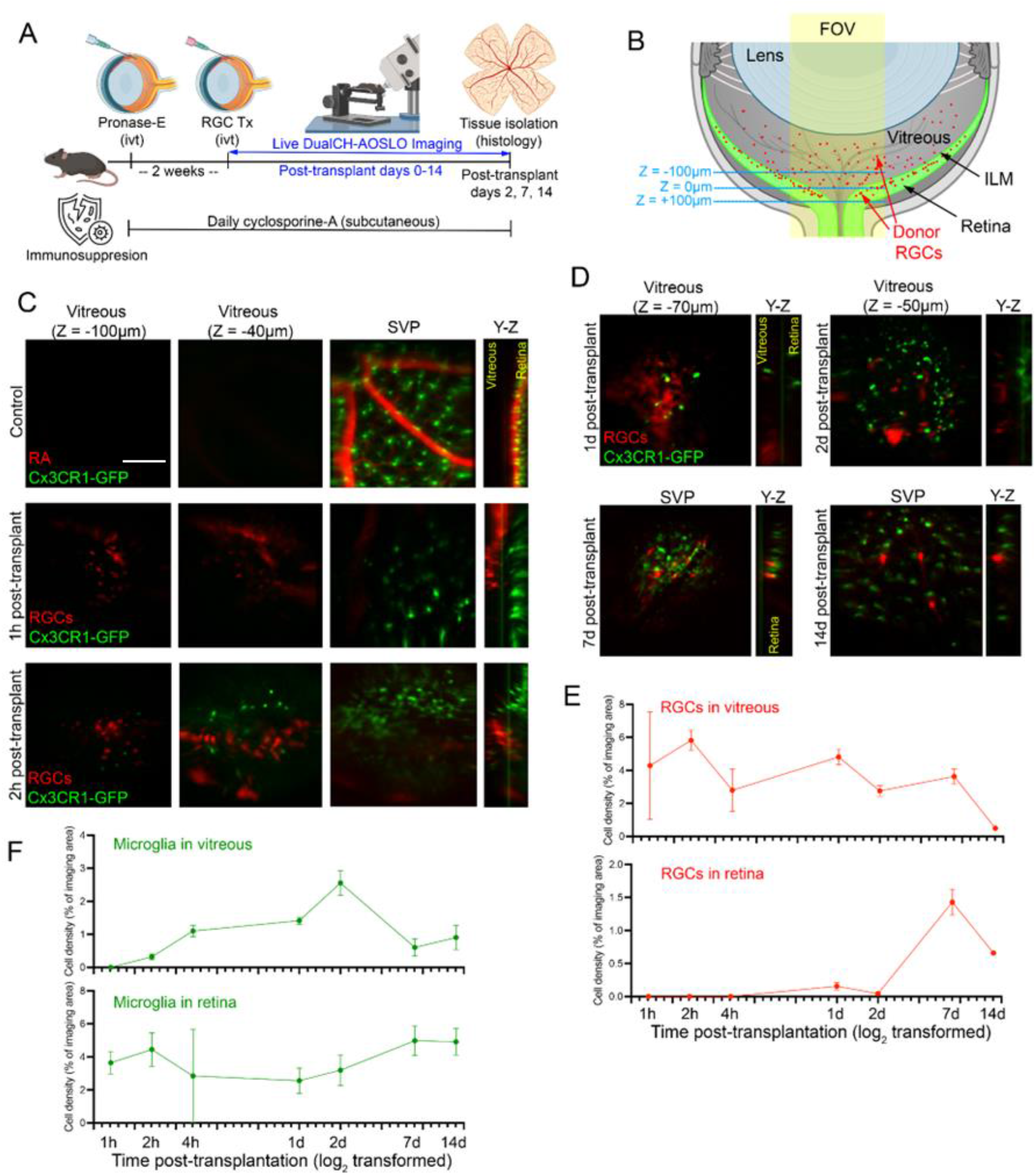
The *in-vivo* 3D imaging of the tRGCs. (A) The schematic of RGC transplantation in CX3CXR-GFP mice. (B) The schematic of imaging locations in RGC transplantation. (C) The pre-IVT retinal imaging of CX3CXR-GFP mice and the early stage of microglia trajectory to the tRGC at 1.5 h and 2.5 h post-transplantation. (D) Long-term in-vivo imaging of tRGC with immune cells on day 1, 2, 7 and 14 post-transplantations. (E) and (F) The cell density (% of imaging area) of RGCs and microglia at different imaging time points in the vitreous and retina. Scale bar: 100 μm.

Prior to or without transplantation, microglia were distributed within the retinal parenchyma with minimal presence in the vitreous cavity (Fig 4C). At 1h post-transplantation, when donor RGCs remained in the vitreous cavity, microglia remained confined to their native retinal layers (Fig 4C). By 2h, microglia began migrating out of the retina and into the vitreous. Over days 1 and 2, donor RGCs remained in the vitreous where microglial engagement increased, characterized by direct contact with donor RGC neurites and cell bodies (Fig 4D). By day 7 and persisting through day 14, donor RGCs had migrated into the inner retina (at the level of the SVP) (Fig. 4D). We quantified RGC and microglial populations across multiple time points by random sampling and calculating the percentage of imaged area that was occupied by each cell type. In the vitreous, transplanted RGCs were maximally populated immediately after transplantation and slowly declined over time (Fig. 4E), In contrast, vitreous microglial infiltration began at 2h and modestly increased through day 2 before subsequently declining (Fig. 4F). In contrast, donor RGCs did not reach the retina until day 7 and they decreased in density of the subsequent week whereas microglial density in the retina remained stable over the 14-day observation period (Fig. 4E, F).

Given the proximity between resident microglia and donor RGCs at early time points when RGC density was dropping significantly, we further examined the morphological relationship between these two cell types using local AO correction to improve the resolution. Static images revealed extensive co-localization between microglial process and donor RGC somas and neurites (Fig 5). Time-lapse imaging over the course of 30-40 minutes revealed examples motile microglia actively extending and remodeling their processes on and around donor RGCs (Fig. 5A), followed shortly after by loss of RGC neurites and disappearance of tdTomato signal from entire neurons (Fig 5A, Supplementary video 8), suggestive of active neurite pruning and death of donor RGCs induced by host microglia.

**Figure 5.**
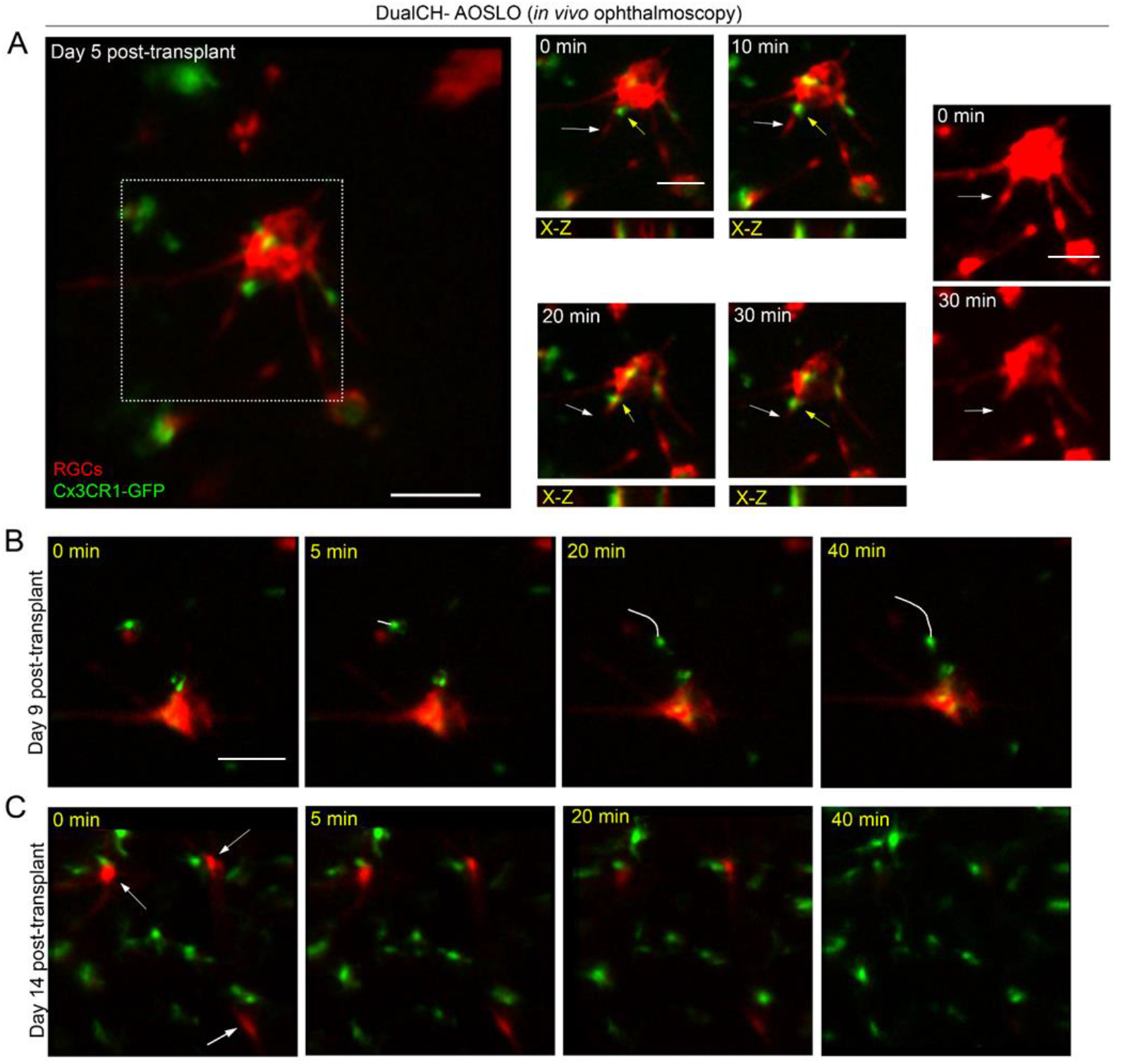
DualCH-AOSLO enables longitudinal in vivo visualization of microglial migration and recruitment toward transplanted RGCs. (A) In vivo 3D AOSLO imaging showing interactions between transplanted RGCs (red) and Cx3cr1-GFP–labeled microglia (green) following transplantation. Time-lapse images and X–Z views reveal dynamic microglial soma and process movements near grafted cells. (B) In vivo time-lapse AOSLO imaging (0–40 min) at day 9 post-transplantation showing directed migration of microglia toward a transplanted RGC soma. The trajectory of a representative migrating microglial cell is indicated. (C) In vivo time-lapse imaging at day 14 post-transplantation showing increased numbers of microglia surrounding transplanted RGCs, suggesting recruitment of microglia to the transplantation site. Scale bar: 100 μm.

As shown in Fig. 5B (Supplementary video 9), microglia exhibited directed migration toward transplanted RGCs, with individual cells progressively moving closer to the graft over a 40-min imaging period. The white trajectory line indicates the migration path of a representative microglial cell. By day 14 post-transplantation, increased numbers of microglia were observed surrounding transplanted RGCs (Fig. 5C, Supplementary video 10), suggesting recruitment of microglia to the transplantation site and ongoing interactions with grafted cells.

Further analysis revealed heterogeneous microglial behaviors within the region. While a subset of microglia exhibited active migration toward transplanted RGCs, others remained relatively stationary. Representative examples of migratory and still microglia are shown in Fig. S7A,B. Quantitative tracking of microglial soma displacement demonstrated that actively migrating microglia displayed significantly higher migration speeds than relatively stationary cells (Fig. S7C).

We validated these *in vivo* imaging findings by directly comparing them to fixed vitreoretinal flatmounts imaged with confocal microscopy (Fig 6A and B). Following transplantation, tdTomato-labeled donor RGCs were detected throughout the preparations, though it was not possible to differentiate RGCs in the vitreous cavity from those on the epiretinal surface due to tissue compression during slide mounting. At early time points (days 2-5), these cells were frequently found in close proximity GFP^+^ microglia. Microglia that clustered around donor RGCs tended to exhibit an activated, ameboid morphology with truncated microglial processes extending toward, contacting, and enwrapping donor RGCs. By days 7-14, fewer donor RGCs remained and some appeared fragmented with punctated tdTomato labeling. Remarkably, at microglia were found across all time points to have engulfed tdT^+^ fragments of donor RGCs, and in some cases entire cells (Fig 6 C,D).

**Fig. 6.**
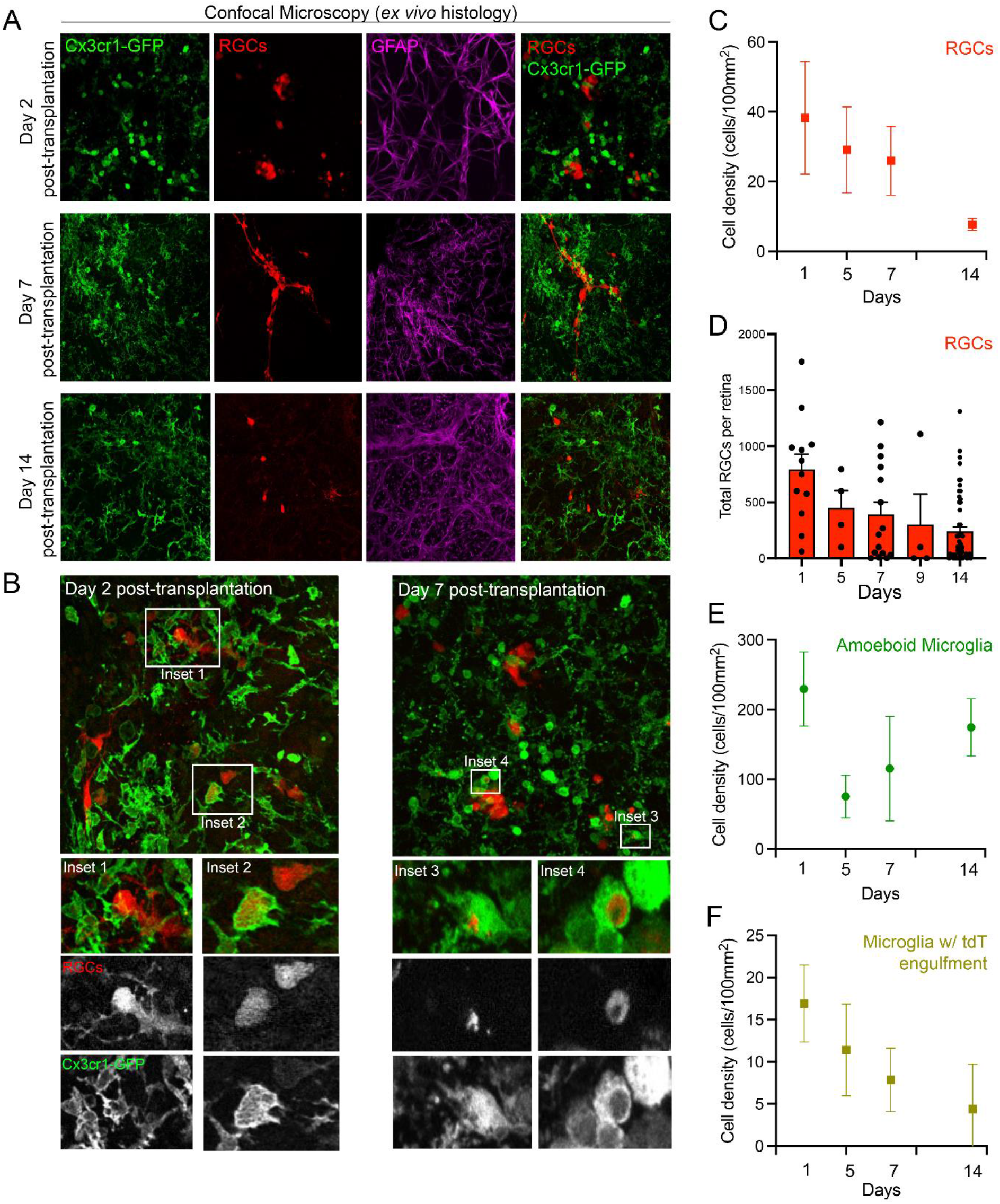
Host microglial response and donor RGC interactions following transplantation. (A) Confocal images of retinal wholemounts at 2, 7, and 14 d post transplantation. Donor retinal ganglion cells (RGCs) expressing tdTomato are shown in red, host microglia in green (Cx3cr1-GFP), host astrocytes in magenta (GFAP), and nuclei are labeled with DAPI. Merged images show the spatial relationship between donor RGCs and host glial cells over time. (B) Higher-magnification images at Days 2 and Days 7 post transplantation highlighting interactions between tdTomato^+^ donor RGCs and host microglia. Insets show representative examples of microglia contacting or engulfing donor-derived material. Individual channels are shown below. (C) Quantification of tdTomato^+^ donor RGC density (cells per 100 mm^2^) over time. (D) Total number of tdTomato^+^ donor RGCs per retina at each time point. Points represent individual retinas; bars indicate mean ± SEM. (E) Density of amoeboid microglia (cells per 100 mm^2^) over time. (F) Density of microglia containing tdTomato^+^ material (cells per 100 mm^2^), indicating phagocytic activity. Scale bar 50 μm.

To further characterize these interactions we quantified the density of GFP^+^ microglia, tdT^+^ donor RGCs, and tdT^+^GFP^+^ microglia that had engulfed donor RGC material or entire cells within randomly sampled fields that contained donor neurons (Fig. 6C-F). Though ameboid (activated) microglial density varied with no clear temporal trajectory (Fig 6E), donor RGC density declined with time (Fig 6C). Consistent with the cell density analysis, quantification of total (non-sampled) donor RGC counts per retina demonstrated a progressive decline over time, with the greatest drop between days 2-5 (Fig. 6D, Kruskal–Wallis p = 0.0024). We then quantified the density of GFP^+^ microglia that had internalized fragments of tdT^+^ RGCs or entire cells (Fig 6B,C). The presence of these phagocytic microglia amounted to approximately half that of the surviving donor RGC density at each time point, following a temporal trajectory that mirrored overall RGC survival (Fig 6F). Together, these findings demonstrate that donor RGC transplantation elicits a robust host microglial response that varies over time and progressively attenuates donor RGC survival.

## Discussion

Recent advances in lens-based AOSLO systems have demonstrated their potential as compact alternatives to traditional mirror-based designs (18, 19). Building on these developments, we present a compact, lens-based DualCH-AOSLO system capable of simultaneous volumetric (3D) two-color fluorescence imaging in the mouse retina. By incorporating polarization optics—including a polarizer and quarter-wave plates—the system effectively suppresses lens surface reflections and minimizes interference with wavefront sensing. Validation using fluorescent microspheres demonstrated near–diffraction-limited transverse and axial resolutions in the red channel, while slightly reduced performance in the green channel suggests residual chromatic effects when using 488-nm excitation for wavefront sensing (see Materials and Methods). Nevertheless, adaptive optics correction substantially enhanced image contrast and signal-to-noise ratio across imaging modalities, particularly in fluorescence angiography. To explore the potential experimental utility of this new technology, we studied microglial dynamics following optic nerve crush and RGC transplantation. DualCH-AOSLO enabled identification of differences in microglial density, morphology, and response to RGC axonal injury according to retinal layer and provided direct evidence of early microglial attack of donor RGCs leading to graft attrition in a transplantation model.

A major limitation of existing AOSLO implementations in mouse models is the restricted usable field of view. Although scanning angle adjustments can extend the nominal FOV to ~10°, diffraction-limited performance is typically confined to much smaller regions (37–39). Even prior lens-based AOSLO systems have largely operated within 3–10° FOVs (20). In contrast, our system achieves a substantially larger AO-corrected FOV (~16°), resulting in more than a twofold improvement in signal-to-noise ratio. This wide-field capability is particularly advantageous for biological studies, as it enables efficient surveying of retinal regions to identify targets of interest before transitioning to high-resolution imaging.

Depth-resolved retinal imaging presents an additional challenge for AOSLO systems. Many previous approaches have relied on electrically tunable lenses (ETLs) to shift the focal plane axially, primarily due to limited deformable mirror (DM) stroke ranges (18, 19). However, ETLs can introduce thermal instability and higher-order aberrations (40). In contrast, the deformable mirror used here provides a substantially larger defocus stroke, sufficient to span the full depth range encompassing the three retinal vascular plexuses (SVP, IVP, and DVP) in the mouse retina. Optical simulations and experimental angiography confirmed a near-linear relationship between DM defocus and retinal imaging depth, enabling coverage of all three vascular plexuses using less than half of the available stroke. Adaptive optics correction markedly improved contrast and axial separation of the SVP, IVP, and DVP, allowing resolution of closely spaced capillaries that were indistinguishable without AO.

These technical advances enabled detailed biological investigations that are difficult to achieve with conventional retinal imaging. We first applied the system to visualize microglia, the primary immune cells of the retina, whose morphology and dynamics are closely linked to inflammatory and degenerative disease states (4, 41, 42). Using depth-resolved reference angiography, we identified distinct microglial populations across retinal layers, with more ramified cells predominantly in the IPL and OPL (corresponding to the IVP and DVP) and fewer microglia in the NFL (SVP), consistent with prior reports (4, 41–43). Simultaneous two-color imaging further demonstrated that AO correction improved image quality in both channels and enabled accurate alignment of vascular and microglial layers in three dimensions, suggesting partial mitigation of chromatic aberration effects.

Beyond static morphology, our system enabled high-resolution, real-time 3D imaging of microglial dynamics. The enhanced resolution revealed fine microglial processes and subtle morphological changes that are critical for understanding immune responses to neurodegenerative insult (44, 45). Quantitative analysis showed microglial process extension and retraction velocities consistent with previous reports (30, 46) and revealed layer-dependent differences in motility, with microglia in the SVP exhibiting greater dynamic activity than those in deeper plexuses in healthy retinas.

To further demonstrate the biological utility of large-FOV, depth-resolved, and dynamic imaging, we applied the system to the optic nerve crush (ONC) model, a widely used paradigm for studying optic neuropathy and axonal injury (4, 47). While ex vivo histology provides high-resolution snapshots of retinal pathology, it lacks the ability to capture temporal evolution in vivo (42, 48). Using our AOSLO platform, we monitored microglial responses longitudinally after ONC. Although gross morphological changes were minimal during the first three days, microglial density increased markedly. By day 8, microglia adopted a radial morphology consistent with activation, accompanied by reduced processes spanning from the SVP to the DVP, indicating heightened immune engagement. These changes persisted through day 21 and were corroborated by ex-vivo confocal imaging, underscoring the value of in-vivo longitudinal observation.

Finally, we leveraged the system’s volumetric and time-resolved capabilities to evaluate host neuroimmune responses to transplanted RGCs. The timing and nature of microglial recruitment to transplanted RGCs aligned with established microglial response kinetics, characterized by rapid morphological change and directed migration within hours after retinal perturbation (4, 48–50). The host microglia underwent a temporally regulated activation following RGC transplantation, engaging with donor neurons early after engraftment and declining as donor cell survival decreases. The peak in microglial density and colocalization at Day 7 suggests an active innate host response phase that may contribute to graft remodeling or clearance, consistent with previous findings that host myeloid populations become activated following neuronal transplantation (35, 43). Early imaging revealed initial migration of microglia out of the retina and into the vitreous with RGC contact within 2 hours, followed by dense clustering around the grafted cell bolus during days 1–2 when transplanted RGCs remained within the vitreous cavity. This very rapid induction of microglial response to donor RGCs suggests that transplantation protocols may benefit from pretreatment with agents that modulate neuroinflammation prior to transplantation. At later stages, transplanted cells transitioned into the superficial vascular plexus, coinciding with a shift toward more dispersed, non-activated microglial morphologies distal to donor RGCs but retention of activated ameboid microglia proximal to and in contact with donor RGCs. Volumetric 3D time-lapse imaging further revealed subacute microglia–neurite interactions, including localized neurite retraction and transient increases in microglial motility, before settling into a late-stage quiescent surveillance state.

Notably, host microglia have been shown in other retinal transplant models to interact closely with donor cells and influence survival outcomes, possibly through phagocytic or trophic mechanisms (35, 51–53). The coincident decline in both microglial engagement and donor RGC counts by Day 14 underscores the importance of the early post-transplant period in determining long-term graft survival, as has been reported in stressed photoreceptors and injured retina (54–58). Future strategies that modulate host microglial activation may improve survival and integration of transplanted neurons, echoing broader themes in neural transplantation research (2, 59–63). Microglia rapidly respond to transplanted neurons, engaging in surveillance, remodeling, and clearance (64, 65). Here, we find that the proportional distribution of microglial–donor RGC interactions remain largely unchanged from D2 to D14. These data suggest that microglial engagement is established early after transplantation and persists rather than progressively escalating over time. Though we observed evidence of microglial internalization of donor RGC derived material, studies incorporating lysosomal markers will be required to determine whether these interactions culminate in functional phagocytosis. Crucially, the timecourse of these interactions emerged only through the combination of large field-of-view coverage, true three-dimensional imaging, and time-resolved observation, highlighting the unique value of this DualCH-AOSLO platform for bridging the gap between static histology and dynamic *in vivo* neuroimmune processes. Ultimately, our data demonstrate that microglial interactions with donor RGCs occur rapidly following transplantation and likely contribute to rapid attrition in graft survival (64, 65).

## Conclusion

In conclusion, we developed a compact, lens-based large-FOV DualCH-AOSLO system that achieves high-resolution, depth-resolved, and temporally stable fluorescence imaging in the mouse retina. By combining wide-field uniformity with dual-color and real-time volumetric capabilities, the system overcomes long-standing limitations of conventional mirror-based AOSLO designs and enables continuous visualization of retinal structures and immune cell behavior with unprecedented fidelity. Using this platform, we revealed previously inaccessible aspects of microglial physiology and neuroimmune interactions. In healthy retina, the system resolved layer-specific microglial dynamics and subtle variations in process motility across vascular plexuses. Following optic nerve crush, it enabled longitudinal tracking of microglial morphological transitions and density changes in vivo, providing insights that are difficult to obtain from single-time-point histology. Most importantly, the system allowed us to delineate a multi-phase microglial response to transplanted RGCs— comprising rapid recruitment, spatial consolidation, and prolonged remodeling—highlighting a sustained, destructive surveillance process that shapes graft–host interactions. Microglial engagement with donor RGCs is strongest during the first week after transplantation. By two weeks, both microglial activity and donor cell numbers have decreased, suggesting that early host responses play a key role in shaping graft survival. Together, these findings demonstrate that wide-field, high-resolution, and time-resolved AOSLO offers a powerful and versatile framework for advancing retinal biology. This technology establishes a foundation for future studies in neuronal repair, immune modulation, and cellular therapy integration, and provides a methodological platform broadly applicable to in vivo neuroimaging and therapeutic monitoring.

## Materials and Methods

### Lens-based optical design

The schematic of the dual-channel adaptive optics scanning laser ophthalmoscope (DualCH-AOSLO) system is illustrated in Fig. 1a and Supp Fig. S9. Briefly, two laser lines (Coherent, 488nm and 552nm) are combined and expanded using a dichroic mirror (DC) and a pair of achromatic relay lenses (L1 and L2) with focal lengths of 25 mm and 150 mm. After passing through a polarizing beamsplitter (PBS), the combined laser beams are expanded threefold again to slightly overfill the aperture of the deformable mirror (DM, ALPAO, DM97-15). The laser beam reflected by the DM is demagnified twofold by another pair of achromatic relay lenses (L6 and L7) with focal lengths of 400 mm and 200 mm, conjugating the DM to the tunable lens (TL: ML-20-37, Optotune Switzerland AG). The beam is then further demagnified by lenses L8 and L9 (focal lengths: 150 mm and 100 mm) to match the 10-mm galvanometer scan mirrors and relay TL to the fast galvanometer mirror. Two slow galvanometer mirrors (GM1 and GM2) (Thorlabs Inc, 6215H), operating with opposite voltage polarity, project the incident beam to focus on the fast-scanning galvanometer mirror (GM3) (66). Finally, the fast-scanning mirror is conjugated to the mouse cornea using a pair of lenses (L10 and L11) with focal lengths of 100 mm and 50 mm, compressing the excitation beam to approximately 2.15 mm in diameter. The entire AOSLO system was installed on a 600 mm-by-600mm breadboard. The detailed layout of the optical system and a photograph are provided in the Supp Figs. S9a and S9b.

For fluorescence detection, a 105 μm core multimode optical fiber is used as the pinhole to guide light to the PMT, The equivalent pinhole size is 5.69 times the air disk diameter (ADD) for mouse retinal imaging for the green channel and 4.85 for the red channel. The wavefront sensor is placed behind the PBS to collect the light returning from retina. In order to remove surface reflections from all the lenses, a quarter wave plate (QWP) was placed after the final lens before the mouse eye (20).

The acquisition software was developed in C++ using the Qt platform (Qt Group Inc., USA) for data collection and AO calculations. During data acquisition, fluorescence signals were collected by the PMT and received by the digitizer (ATS9440, Alazar Technologies Inc.). The raw data was then processed and displayed in real-time by the acquisition software. Additionally, our software controls the synchronization of the galvo scanners and the digitizer. For AO calculations and closed-loop operation, the acquisition software integrates the MATLAB Engine to control the WFS and DM.

### Validation of AO correction and simultaneous 3D in-vivo imaging of GFP-labeled microglia and rhodamine angiography

To validate the performance of AO correction, a 3-mm N-BK7 ball lens (Edmund Optics, #43-711) was used to emulate the mouse eye with comparable size and numerical aperture (NA) (Supp Fig. S10). A diluted suspension of yellow-green (A803431, Polysciences Inc.) and red fluorescent microspheres (A800977, Polysciences Inc.), each with a diameter of 0.5 μm, was prepared. A small droplet of the mixed solution was deposited onto lens cleaning tissue and allowed to dry. To optimize system resolution, the multimode fiber coupled to the photomultiplier tube (PMT) was replaced with a 25 μm-core fiber, yielding effective ADD values of 1.44 for the green channel and 1.9 for the red channel.

### Simulation of imaging plane with changes of DM defocus term

To explore the relationship between the DM defocus term and imaging plane shift in the retina or other objects, optical models using a 3 mm N-BK7 glass ball and a mouse eye model (67) were designed and simulated in Zemax OpticStudio (Ansys, Inc.), as shown in Supp Fig. S10. A mathematical model was developed to examine the relationship between the radius *R*_*WF*_ of the DM surface arc and the DM surface shift *Δd* resulting from the defocus Zernike term (Supp Fig. S11A).

The wavefront curvature *R*_WF_was incorporated into the Zemax model by modifying the deformable mirror (DM) surface, as illustrated in Supp Fig. S11B C. By varying *R*_WF_, the corresponding shifts of the focal plane were recorded. The relationship between the focal plane shift and Δ*d*was then established (Equation 3), revealing a linear correlation between these two parameters, as confirmed by the simulation results shown in Supp Fig. S11c.

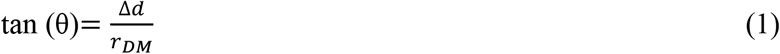

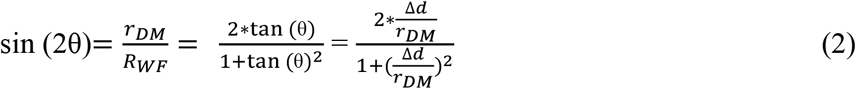

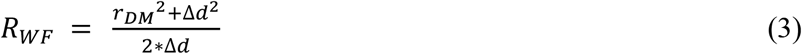

### Animal preparation and imaging

All experiments were conducted in accordance with the ethical guidelines for animal care and received approval from the Institutional Animal Care and Use Committee at Johns Hopkins University (Protocol#M021M350). Two C57BL/6J wild-type mice and 45 *Cx3Cr-1GFP* mice were used in this study. Throughout the imaging process, a mixture of 1% (v/v) isoflurane and oxygen was administered to maintain stable anesthesia, while a heat pad kept the body temperature at 37°C. After the mice were dilated and anesthetized with ketamine/xylazine, 0.1 ml of 2.5% Fluorescein isothiocyanate–dextran (46944-500MG-F, Sigma-Aldrich Co) or 0.1 ml of 2.5% Rhodamine B isothiocyanate–Dextran (R9379-100MG, Sigma-Aldrich Co) was injected retro-orbitally to enable visualization of the retinal vasculature. A +10 D contact lens with hypromellose gel was applied to the cornea to prevent dehydration and cataract formation during imaging.

Although the system supports imaging up to ~20°, most AO experiments were performed using ~16° sensing regions to maintain stable wavefront estimation and uniform image quality. The mice were then positioned on the imaging stage for *in-vivo* retinal imaging using the DualCH-AOSLO. Similar procedures were followed for CX3CR1-GFP mice, but only Rhodamine-B dextran was used for angiography. For AO imaging of fluorescein or GFP, the 488 nm laser was activated for wavefront sensing (552 nm laser for the wavefront sensing during rhodamine angiography), focusing on imaging planes around the inner vascular plexus (IVP) layers by adjusting the TL. The AO closed loop was then initiated to correct for aberrations.

For simultaneous 3D dual-channel data acquisition, both the 488 nm and 552 nm lasers were activated, and a 512 × 512 × 100 image stack was recorded. To study the dynamics of microglial cells, imaging fields containing 1 to 4 cells in the superficial vascular plexus (SVP) layers were selected, and TL was adjusted to confirm the morphology of microglial cells in the IVP and deep vascular plexus (DVP). During imaging, Rhodamine-B angiography was used to maintain the field of view on the same region of interest, ensuring consistent imaging locations throughout the time-lapse imaging.

To study the dynamics of microglial processes, 0.1 mW of 488 nm laser was used, and the focus plane was shifted to the SVP by adjusting the TL. After opening the AO closed loop and ensuring the Strehl ratio of the wavefront map exceeded 0.8, a 512 × 512 × 50 imaging stack was collected at each time point. Dynamic imaging was repeated immediately after data acquisition and every 5 minutes thereafter. Each dynamic imaging session typically lasted between 35 to 95 minutes for each CX3CR1 mouse, depending on image quality, depth of anesthesia, and animal cooperation.

### Optic nerve crush

ONC surgery was conducted following the procedures described previously (68, 69). Briefly, mice were anesthetized with isoflurane (as above) and the ocular surface was prepped with 0.5% proparacaine and 5% betadine. A lateral conjunctival incision was made using Vannas scissors and blunt dissection was performed to visualize the optic nerve, taking care not to disrupt the orbital blood circulation. The optic nerve was crushed for 3 seconds using calibrated cross-action forceps. Confirmation of retinal blood flow was made immediately after through direct visualization of the fundus. After ONC, the mouse was imaged on days 1, 2, 3, 8, 10, 14, and 21. On day 21, mice were sacrificed after *in-vivo* imaging, and the retinas were extracted and processed for confocal immunofluorescent microscopy. *Ex-vivo* imaging was performed using a commercial confocal microscope (Zeiss LSM 880 confocal laser scanning microscope, Carl Zeiss Microscopy LLC, White Plains NY).

#### Human Stem Cell Culture and RGC Differentiation

Human H7 embryonic stem cells (hESCs) engineered to express tdTomato and CD90.2/Thy1.2 at the endogenous POU4F2 (BRN3B) locus (gift from D. Zack, Johns Hopkins University) were maintained on Matrigel in mTeSR1 medium at 37°C, 5% CO_2_ (70). Cells were passaged using Accutase and replated as single-cell suspensions with 5 µM blebbistatin. RGC differentiation was induced using sequential small-molecule treatment in iNS medium over 30 days, including Forskolin, Dorsomorphin, IDE2, DAPT, and Nicotinamide at defined temporal windows(71, 72).

#### RGC Purification and Cryopreservation

On day 40, cells were dissociated with Accumax and purified by anti-CD90.2 MACS (Miltenyi Biotec), achieving 95–99% purity confirmed by flow cytometry(71). Purified RGCs were resuspended in CryoStor CS10 at 4×10^6^ cells/vial, frozen at ™80°C overnight, and transferred to liquid nitrogen. At use, cells were thawed at 37°C, washed in BrainPhys neuronal medium, and counted prior to transplantation.

### Intravitreal Pronase-E and Cell Injections

Prior to cell transplantation, mice received an intravitreal (IVT) injection of Pronase -E (0.14U/mL; Catalog #P8811-1G, Lot #0000355919 cas no: 9036-06-0) LOT specific enzyme activity: 3.5 units/mg to digest the inner limiting membrane and facilitate donor cell integration. Two weeks post Pronase-E treatment, donor RGCs (H7RGC) were prepared, resuspended at a concentration of 37,500 cells/1.5 μL/ injected under sterile conditions.

### RGC transplantation

To minimize immune rejection of transplanted RGCs, mice received subcutaneous injections of cyclosporine A (25 mg/kg/day) at the beginning one week prior to transplantation and continuing until the experimental endpoint.

### Retina dissection and Immunostaining (73)

Mice were humanely euthanized by ketamine/xylazine overdose, followed by cervical dislocation. Under a microscope, a small portion of the eyelid was removed using straight forceps for better visualization. The conjunctiva was held with forceps while micro scissors were used to make small incisions, avoiding damage to the eyeball, optic nerve head, and optic nerve. The extracted eyes were transferred into labeled Falcon tubes containing cold PBS. Eyes were submerged in PBS to facilitate tissue removal and maintain shape.

A syringe needle was used to create a small hole at the cornea-sclera junction, followed by a circumferential incision with micro scissors to remove the cornea and iris. The lens was gently expelled by pressing on the posterior eyecup. The neural retina was detached and incised radially to create a clover-like shape and transferred onto a membrane with the photoreceptor side down. Retinas were placed in 4% PFA for one hour at 4°C. The fixative was then replaced with PBS, and samples were stored at 4°C until further processing.

Fixed retinal explants were washed in PBS (3 × 5 min) while rocking. Retinas were then incubated in blocking solution (1× PBS, 0.3% Triton X-100, 10% normal goat serum) for 60 min at room temperature with gentle rocking. Retinas were incubated with primary antibodies RFP (1:400) -cat # 600-901-379 chicken Rockland 42649 to identify transplanted RGCs, GFP (ck-1:1000 (GFP 1020 chicken Aves GFP917979), RBPMS-rb-1:500 (1830 rabbit Phosphosolutions NB523r / NB1223y / NB923A), GFAP rt-1:2000 (2.2B10) 13-300 rat thermos VB298933/ XD346000), diluted in blocking solution (300 µL) for five overnights at 4°C in the dark with rocking. After primary incubation, retinas were washed in PBS (3 × 5 min) and blocked again for 60 min at room temperature. Secondary antibodies Goat anti Chicken Alexa-488 cat# A11039 thermo-lot# 2304258/ 2566343, GoatαRabbit Alexa-568cat # A11011 GoatαRabbit thermo-lot # 2500544 (2X), Goat-anti-Rat-Alexa-647 cat # A21247 thermo-lot# 2519439 were applied in blocking solution (500 µL) and incubated overnight at 4°C in the dark with gentle rocking. Retinas were washed in PBS (3 × 5 min), incubated in DAPI 4′,6-Diamidine-2′-phenylindole dihydrochloride cat#-10236276001 Millipore Sigma for nuclear counterstaining (1:2000 in PBS) for 15 min while rocking, and washed again in PBS for 15 min. Retinas were then transferred onto labeled slides using forceps under a microscope, cover slipped with hard-set mounting media -Aqua-poly/mount-18606-5 from poly science, and stored at 4°C in the dark until imaging. 40X high resolution Imaging was done on the 880 confocal microscope (Zeiss LSM 880 confocal laser scanning microscope, Carl Zeiss Microscopy LLC, White Plains NY.) and olympus Fluoview 4000.

### Imaging processing and registration

All images were processed and registered using ImageJ (NIH) and MATLAB (MathWorks Inc). Typically, most volumetric (3D) image stacks exhibited shift artifacts along the Z-axis due to breathing or positional drift of the mouse holder. To eliminate motion-induced artifacts, image registration using a demon-based non-rigid registration algorithm was performed in MATLAB. Subsequently, the image data was smoothed using a Gaussian filter (kernel size: 5×5 pixels).

### Statistics

To quantitatively analyze the dynamics of microglia in different retinal layers, 32 microglial cells from six CX3CR1-GFP healthy mice were selected for analysis. This included 9 cells from the SVP layer, 12 cells from the IVP layer, and 11 cells from the DVP layer. The correlations at each time point for each microglial cell, relative to the initial imaging point (0 min), were used for quantitative comparisons among the three retinal layers. Differences between groups were analyzed using unpaired one-way ANOVA tests in SPSS® 29 software (IBM Inc.). A p-value < 0.05 (*) and p-value < 0.005 (**) were considered statistically significant.

### Data Availability

Raw and processed imaging data will be made available upon reasonable request. BioRxiv DOI: 10.1101/2025.03.31.645335.

### Code Availability

Custom MATLAB code for image reconstruction and 3D rendering will be provided upon request and may be made public upon publication.

## Acknowledgements

We thank the following funding sources supporting this study: NIH R01EY032163, P30EY001765, K08EY031801, R21EY034332, Wilmer Pooled Professorship funding, Research to Prevent Blindness (Career Development Award), Bright Focus Foundation G2022005S, The Glaucoma Foundation Rajen Savjani Award, The Shelley and Allan Holt Rising Professorship, and The Zenkel Family Foundation. We thank Arumugam Nagalingam for the invaluable assistance with cell culture work in generating stem cell–derived retinal ganglion cells for transplantation.

## Author contributions

**J.Y**. and **T.V.J**. conceived the project and supervised all aspects of the study. **Z.L**. designed and constructed the optical system with input from **J.Y**.. **S.M**. and **Z.L**. performed animal imaging experiments and data acquisition. **Z.L**. developed custom software for image reconstruction and 3D rendering. **T.V.J**. provided biological guidance and performed the surgery for the ONC injury model and interpreted neuroinflammatory findings. **Z.L**. and **S.M**. analyzed the data. **Z.L**. and **S.M**. wrote the manuscript with input from all authors. All authors discussed the results and contributed to the final manuscript.

## Competing interests

All the authors declare no potential conflicts of interest to the authorship and/or publication in this article.

## Materials & Correspondence

The DualCH-AOSLO system was custom-built incorporating polarization-sensitive optics, a high-stroke deformable mirror, and lens-based refractive elements optimized to minimize off-axis aberrations. Detailed system configuration and component specifications are provided in the Methods section and Supplementary Information. Transgenic mice expressing microglial markers were used for longitudinal in vivo imaging to track neuroimmune responses following optic nerve crush injury. All animal experiments were performed in accordance with institutional ethical guidelines.

## Notes

### Competing Interest Statement

The authors have declared no competing interest.

### Summary of Updates

we have added more findings and results

## References

1. T. Y. P. Chui et al., Human retinal microvascular imaging using adaptive optics scanning light ophthalmoscopy. Int J Retina Vitreous 2, 11 (2016).

2. K. Y. Zhang, E. A. Aguzzi, T. V. Johnson, Retinal ganglion cell transplantation: approaches for overcoming challenges to functional integration. Cells 10, 1426 (2021).

3. Y.-R. Wu et al., Transplanted mouse embryonic stem cell–derived retinal ganglion cells integrate and form synapses in a retinal ganglion cell-depleted mouse model. Investigative Ophthalmology & Visual Science 62, 26–26 (2021).

4. N. D. Heuss et al., Optic nerve as a source of activated retinal microglia post-injury. Acta Neuropathol Commun 6, 66 (2018).

5. F. Ginhoux et al., Fate mapping analysis reveals that adult microglia derive from primitive macrophages. Science 330, 841–845 (2010).

6. M. Mujat, R. D. Ferguson, D. X. Hammer, A. H. Patel, N. Iftimia (2024) High-Resolution Retinal Imaging: Technology Overview and Applications. in Photonics (MDPI), p 522.

7. E. Bakker et al., Adaptive optics ophthalmoscopy: a systematic review of vascular biomarkers. Surv Ophthalmol 67, 369–387 (2022).

8. P. Godara, A. M. Dubis, A. Roorda, J. L. Duncan, J. Carroll, Adaptive optics retinal imaging: emerging clinical applications. Optometry and vision science: official publication of the American Academy of Optometry 87, 930 (2010).

9. L. X. Liu et al., Application of Adaptive Optics in Ophthalmology. Photonics 9 (2022).

10. Y. Geng et al., Optical properties of the mouse eye. Biomed Opt Express 2, 717–738 (2011).

11. Y. Geng et al., Adaptive optics retinal imaging in the living mouse eye. Biomed Opt Express 3, 715–734 (2012).

12. D. Scoles et al., In vivo imaging of human cone photoreceptor inner segments. Investigative ophthalmology & visual science 55, 4244–4251 (2014).

13. A. Roorda, Applications of adaptive optics scanning laser ophthalmoscopy. Optom Vis Sci 87, 260–268 (2010).

14. A. Roorda et al., Adaptive optics scanning laser ophthalmoscopy. Optics express 10, 405–412 (2002).

15. L. K. Young, T. J. Morris, C. D. Saunter, H. E. Smithson, Compact, modular and in-plane AOSLO for high-resolution retinal imaging. Biomed Opt Express 9, 4275–4293 (2018).

16. B. Moon et al., Alignment, calibration, and validation of an adaptive optics scanning laser ophthalmoscope for high-resolution human foveal imaging. Appl Opt 63, 730–742 (2024).

17. R. D. Ferguson et al., Adaptive optics scanning laser ophthalmoscope with integrated wide-field retinal imaging and tracking. JOSA A 27, A265–A277 (2010).

18. Y. Yang et al., Retinal microvascular and neuronal pathologies probed in vivo by adaptive optical two-photon fluorescence microscopy. eLife 12 (2023).

19. Z. Qin et al., Adaptive optics two-photon microscopy enables near-diffraction-limited and functional retinal imaging in vivo. Light Sci Appl 9, 79 (2020).

20. F. Felberer, J. S. Kroisamer, C. K. Hitzenberger, M. Pircher, Lens based adaptive optics scanning laser ophthalmoscope. Opt Express 20, 17297–17310 (2012).

21. T. DuBose et al., Handheld Adaptive Optics Scanning Laser Ophthalmoscope. Optica 5, 1027–1036 (2018).

22. M. J. Ju, C. Huang, D. J. Wahl, Y. Jian, M. V. Sarunic, Visible light sensorless adaptive optics for retinal structure and fluorescence imaging. Optics letters 43, 5162–5165 (2018).

23. J. Z. Liang, D. R. Williams, D. T. Miller, Supernormal vision and high-resolution retinal imaging through adaptive optics. J Opt Soc Am A 14, 2884–2892 (1997).

24. D. Valente, K. V. Vienola, R. J. Zawadzki, R. S. Jonnal, Kilohertz retinal FF-SS-OCT and flood imaging with hardware-based adaptive optics. Biomedical Optics Express 11, 5995–6011 (2020).

25. V. P. Pandiyan et al., High-speed adaptive optics line-scan OCT for cellular-resolution optoretinography. Biomedical optics express 11, 5274–5296 (2020).

26. R. J. Zawadzki et al., Adaptive-optics SLO imaging combined with widefield OCT and SLO enables precise 3D localization of fluorescent cells in the mouse retina. Biomedical optics express 6, 2191–2210 (2015).

27. F. Felberer et al., Adaptive optics SLO/OCT for 3D imaging of human photoreceptors in vivo. Biomedical optics express 5, 439–456 (2014).

28. D. Merino, P. Loza-Alvarez, Adaptive optics scanning laser ophthalmoscope imaging: technology update. Clin Ophthalmol 10, 743–755 (2016).

29. S. A. Mills et al., Microglia regulate vascular structure in the developing retina. Proceedings of the National Academy of Sciences 118, e2112561118 (2021).

30. A. Joseph, D. Power, J. Schallek, Imaging the dynamics of individual processes of microglia in the living retina in vivo. Biomed Opt Express 12, 6157–6183 (2021).

31. E. G. O’Koren et al., Microglial function is distinct in different anatomical locations during retinal homeostasis and degeneration. Immunity 50, 723-737. e727 (2019).

32. J. Singaravelu, L. Zhao, R. N. Fariss, W. T. Wong, Microglia in the primate retina: regional distribution and morphological diversity. Brain Structure and Function 222, 2759–2771 (2017).

33. M. R. Damani et al., Age-related alterations in the dynamic behavior of microglia. Aging Cell 10, 263–276 (2011).

34. J. Oswald, E. Kegeles, T. Minelli, P. Volchkov, P. Baranov, Transplantation of miPSC/mESC-derived retinal ganglion cells into healthy and glaucomatous retinas. Molecular Therapy-Methods & Clinical Development 21, 180–198 (2021).

35. E. Kriukov et al., Single-cell transcriptome of retinal myeloid cells in response to transplantation of human neurons reveals reversibility of microglial activation. bioRxiv (2025).

36. E. A. Aguzzi et al., Internal limiting basement membrane inhibits functional engraftment of transplanted human retinal ganglion cells. Science Translational medicine (2026).

37. P. Zhang et al., Adaptive optics scanning laser ophthalmoscopy and optical coherence tomography (AO-SLO-OCT) system for in vivo mouse retina imaging. Biomed Opt Express 14, 299–314 (2023).

38. P. Zhang, M. Goswami, A. Zam, E. N. Pugh, R. J. Zawadzki, Effect of scanning beam size on the lateral resolution of mouse retinal imaging with SLO. Opt Lett 40, 5830–5833 (2015).

39. H. Terasaki, S. Sonoda, M. Tomita, T. Sakamoto, Recent Advances and Clinical Application of Color Scanning Laser Ophthalmoscope. J Clin Med 10 (2021).

40. I. Grulkowski et al., Swept source optical coherence tomography and tunable lens technology for comprehensive imaging and biometry of the whole eye. Optica 5 (2018).

41. Y. Liu et al., Monitoring retinal morphologic and functional changes in mice following optic nerve crush. Invest Ophthalmol Vis Sci 55, 3766–3774 (2014).

42. P. Yang et al., Progesterone alters the activation and typing of the microglia in the optic nerve crush model. Exp Eye Res 212, 108805 (2021).

43. N. P. B. Au, C. H. E. Ma, Neuroinflammation, Microglia and Implications for Retinal Ganglion Cell Survival and Axon Regeneration in Traumatic Optic Neuropathy. Front Immunol 13, 860070 (2022).

44. H. Zhang, C. Tan, X. Shi, J. Xu, Impacts of autofluorescence on fluorescence based techniques to study microglia. BMC Neurosci 23, 21 (2022).

45. N. Stence, M. Waite, M. E. Dailey, Dynamics of microglial activation: a confocal time-lapse analysis in hippocampal slices. Glia 33, 256–266 (2001).

46. A. Nimmerjahn, F. Kirchhoff, F. Helmchen, Resting microglial cells are highly dynamic surveillants of brain parenchyma in vivo. Science 308, 1314–1318 (2005).

47. A. M. Hilla, H. Diekmann, D. Fischer, Microglia Are Irrelevant for Neuronal Degeneration and Axon Regeneration after Acute Injury. J Neurosci 37, 6113–6124 (2017).

48. S. G. Wohl, C. W. Schmeer, O. W. Witte, S. Isenmann, Proliferative response of microglia and macrophages in the adult mouse eye after optic nerve lesion. Invest Ophthalmol Vis Sci 51, 2686–2696 (2010).

49. S. E. Haynes et al., The P2Y12 receptor regulates microglial activation by extracellular nucleotides. Nature neuroscience 9, 1512–1519 (2006).

50. A. M. Fontainhas et al., Microglial morphology and dynamic behavior is regulated by ionotropic glutamatergic and GABAergic neurotransmission. PloS one 6, e15973 (2011).

51. Q. Ren, F. Lu, R. Hao, Y. Chen, C. Liang, Subretinal microglia support donor photoreceptor survival in rd1 mice. Stem Cell Research & Therapy 15, 436 (2024).

52. V. V. Malechka et al., Improvement of human donor retinal ganglion cell survival through modulation of microglia. bioRxiv, 2024.2007. 2006.602307 (2024).

53. R. Banerjee, R. Lund, A role for microglia in the maintenance of photoreceptors in retinal transplants lacking pigment epithelium. Journal of neurocytology 21, 235–243 (1992).

54. L. Zhao et al., Microglial phagocytosis of living photoreceptors contributes to inherited retinal degeneration. EMBO molecular medicine 7, 1179–1197 (2015).

55. C. M. Eandi et al., Subretinal mononuclear phagocytes induce cone segment loss via IL-1β. Elife 5, e16490 (2016).

56. Y. Okunuki et al., Microglia inhibit photoreceptor cell death and regulate immune cell infiltration in response to retinal detachment. Proceedings of the National Academy of Sciences 115, E6264–E6273 (2018).

57. R. Sudharsan et al., Metabolic stress and early cell death in photoreceptor precursor cells following retinal transplantation. Stem Cell Research & Therapy 16, 397 (2025).

58. M. H. Madeira, R. Boia, P. F. Santos, A. F. Ambrósio, A. R. Santiago, Contribution of microglia-mediated neuroinflammation to retinal degenerative diseases. Mediators of inflammation 2015, 673090 (2015).

59. M. Andreu et al., Dose-dependent modulation of microglia activation in rats after penetrating traumatic brain injury (pTBI) by transplanted human neural stem cells. PLoS One 18, e0285633 (2023).

60. R. A. Barker, M. Götz, M. Parmar, New approaches for brain repair—from rescue to reprogramming. Nature 557, 329–334 (2018).

61. Z. Kokaia, G. Martino, M. Schwartz, O. Lindvall, Cross-talk between neural stem cells and immune cells: the key to better brain repair? Nature neuroscience 15, 1078–1087 (2012).

62. D. Nayak, T. L. Roth, D. B. McGavern, Microglia development and function. Annual review of immunology 32, 367–402 (2014).

63. M. Politis, O. Lindvall, Clinical application of stem cell therapy in Parkinson’s disease. BMC medicine 10, 1 (2012).

64. B. Stevens et al., The classical complement cascade mediates CNS synapse elimination. Cell 131, 1164–1178 (2007).

65. D. P. Schafer et al., Microglia sculpt postnatal neural circuits in an activity and complement-dependent manner. Neuron 74, 691–705 (2012).

66. W. Shao, J. Yi, Non-interferometric volumetric imaging in living human retina by confocal oblique scanning laser ophthalmoscopy. Biomedical Optics Express 13, 3576–3592 (2022).

67. M. R. R. Gardner, H. Grady; Milner, Thomas E. (2016) Computational Model (Zemax) of a Mouse Eye for Visible and Near-Infrared Wavelength Ranges. (The University of Texas at Austin, The University of Texas at Austin).

68. E. G. Cameron et al., Optic Nerve Crush in Mice to Study Retinal Ganglion Cell Survival and Regeneration. Bio Protoc 10 (2020).

69. N. D. Heuss et al., Retinal dendritic cell recruitment, but not function, was inhibited in MyD88 and TRIF deficient mice. Journal of Neuroinflammation 11, 1–16 (2014).

70. J. Maruotti et al., A simple and scalable process for the differentiation of retinal pigment epithelium from human pluripotent stem cells. Stem cells translational medicine 2, 341–354 (2013).

71. V. M. Sluch et al., Enhanced stem cell differentiation and immunopurification of genome engineered human retinal ganglion cells. Stem cells translational medicine 6, 1972–1986 (2017).

72. W. R. Institute (2024) H7 Human Embryonic Stem Cell Line. (WiCell Research Institute, Madison, WI, USA).

73. K. Y. Zhang, T. V. Johnson, Analyses of transplanted human retinal ganglion cell morphology and localization in murine organotypic retinal explant culture. STAR protocols 3, 101328 (2022).

