## Supplemental Fig for "Compact refractive dual-channel AOSLO for wide-field imaging in mice reveals microglial interactions with transplanted neurons"

**Results**

Figure S1:

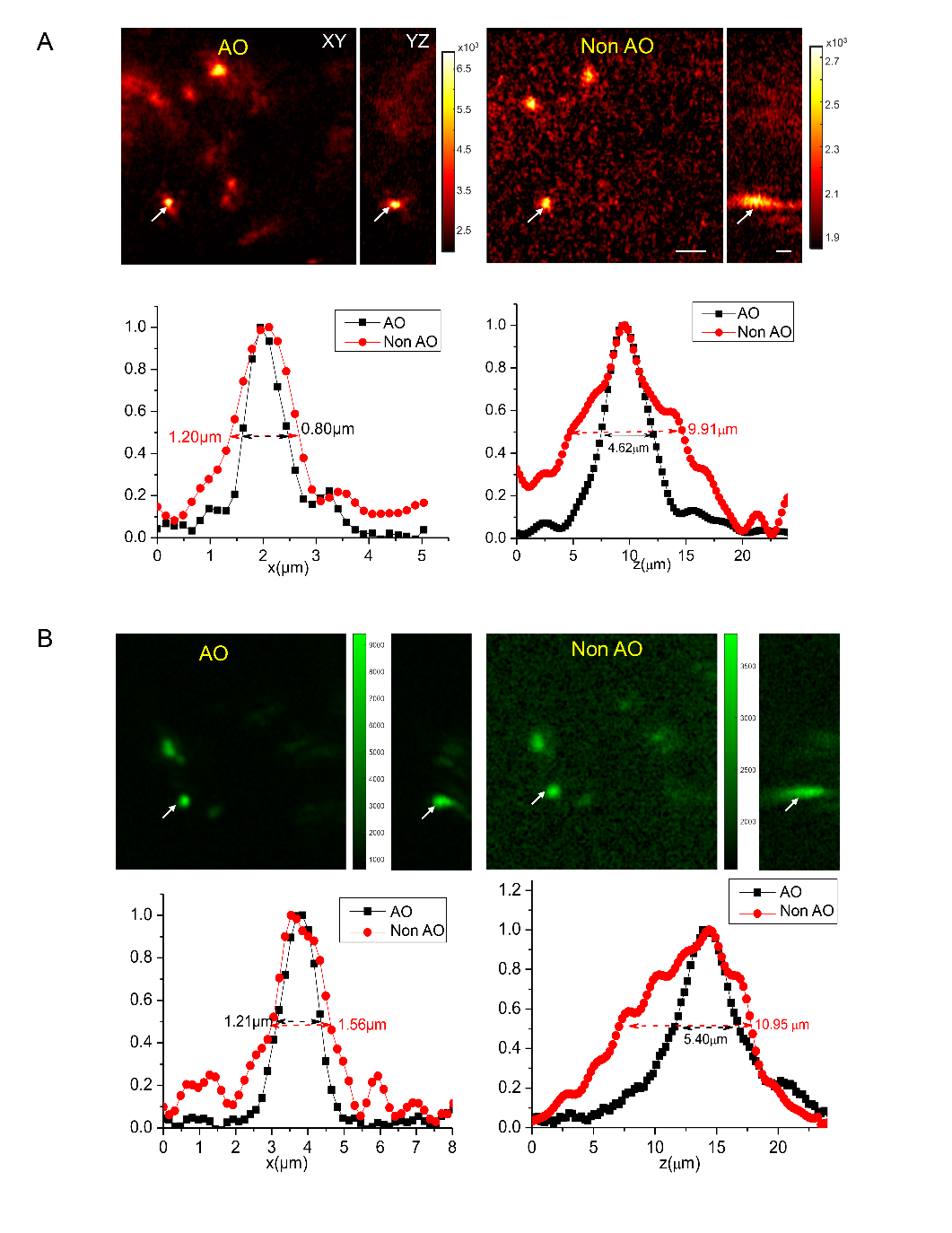

**Figure S1. The imaging resolution with 0.5 μm fluorescence beads (Green).** The imaging resolution with 0.5 μm Red (A) and Green (B) fluorescence beads with AO correction (AO) and without AO correction (NO-AO); The transverse and axial cross section of the Red (A) and Green (B) fluorescence beads that were used to estimate the lateral and axial imaging resolution before and after AO correction.

Figure S2:

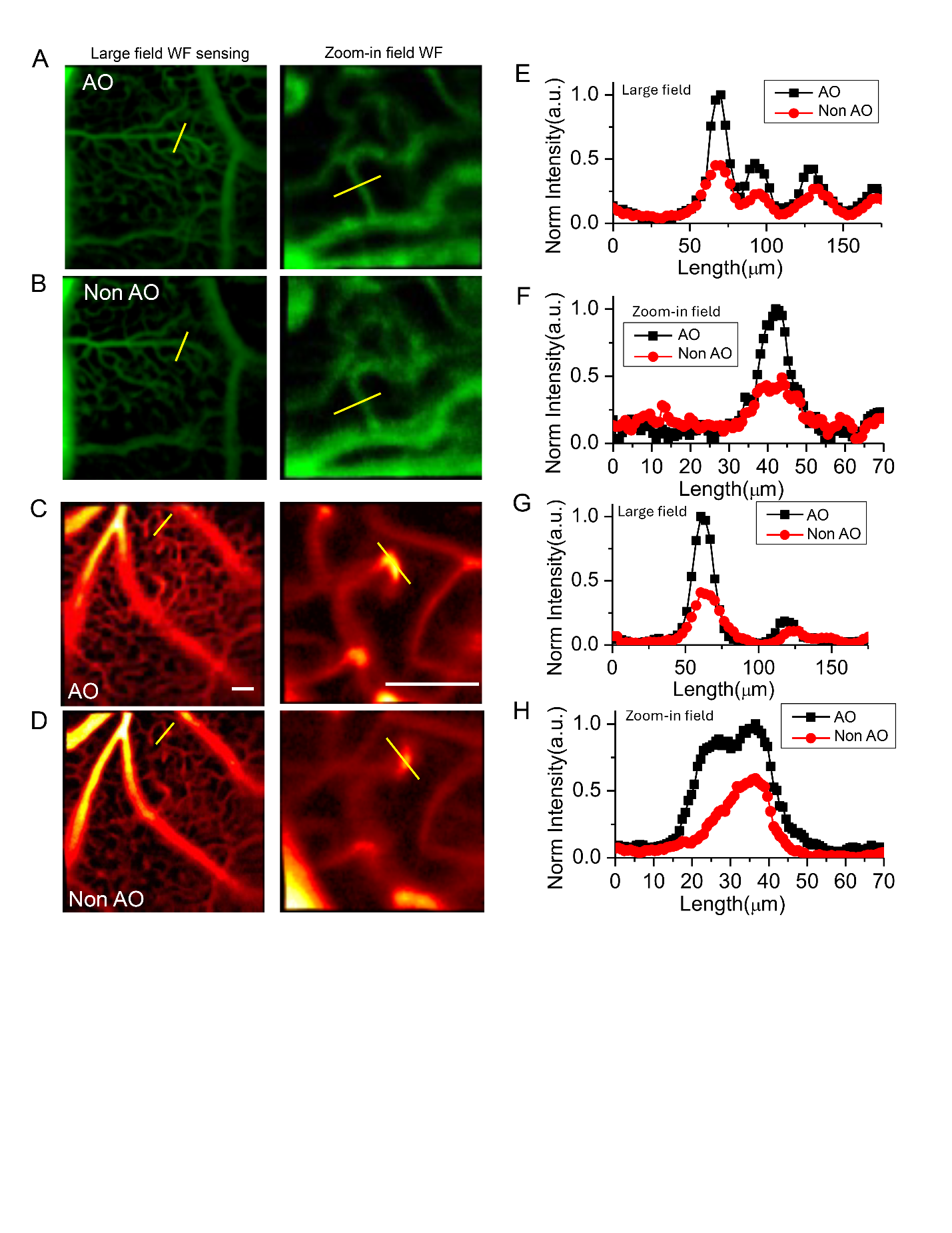

**Figure S2. The RA and FA of DualCH-AOSLO with and without AO correction and their evaluation under large field and small field wavefront sensing.** (A) and (B) The FA angiography with and without AO correction under large and small imaging fields. (C) and (D) The RA angiography with and without AO correction under large and small imaging fields. (E-H) The normalized pixel intensity of AO and NO AO correction under large and small imaging fields in RA.

Figure S3:

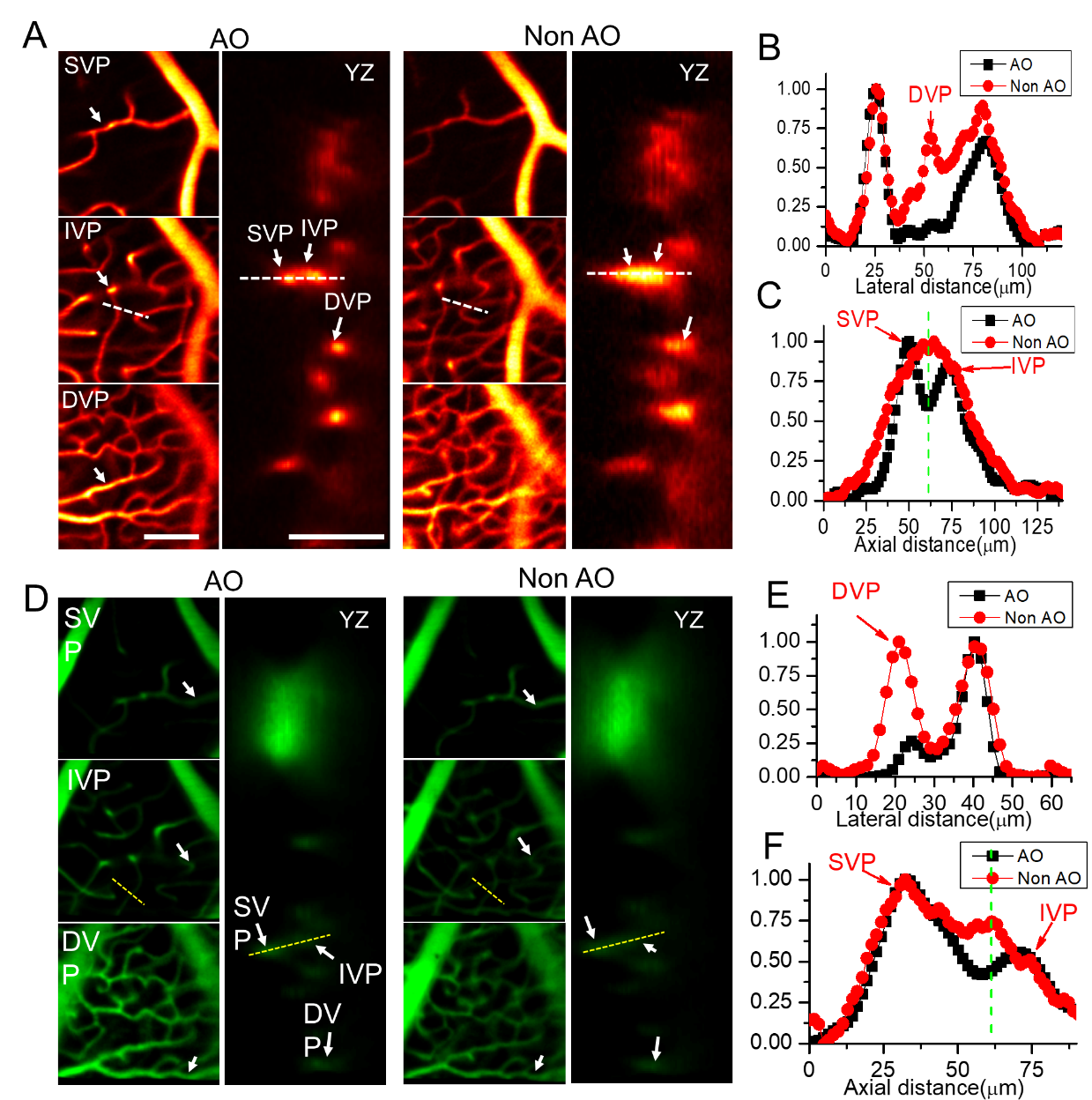

**Figure S3. Characteristics of dual-channel AOSLO for RA and FA imaging under AO and non-AO conditions.** (A) Distinguishing the three retinal vascular plexuses in XY and YZ views for RA imaging. (B–C) Quantitative intensity profile analysis of adjacent capillaries in RA imaging under AO and non-AO conditions, showing improved lateral separation of overlapping DVP signals and enhanced axial discrimination between the SVP and IVP with AO correction. (D) Distinguishing the three retinal vascular plexuses in XY and YZ views for FA imaging. (E–F) Quantitative intensity profile analysis of adjacent capillaries in FA imaging under AO and non-AO conditions, showing improved lateral separation of overlapping DVP signals and enhanced axial discrimination between the SVP and IVP with AO correction. Scale bar: 100 μm.

Figure S4:

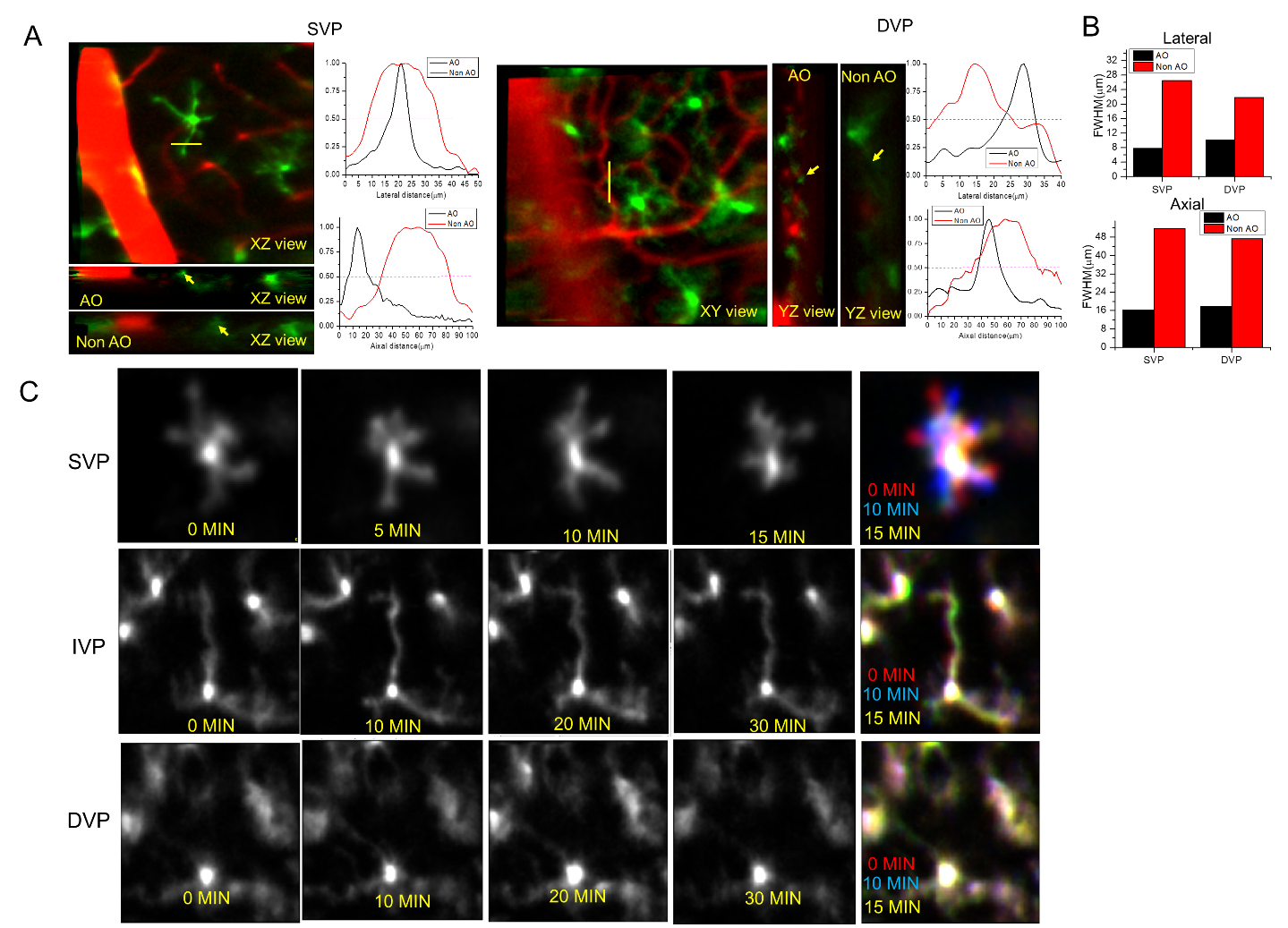

**Figure S4. AO improves the spatial resolution of microglial processes across retinal vascular plexuses, and dynamic of MCs in three retinal layers at different time points.** (A) Representative microglial process images and corresponding lateral and axial intensity profiles in the SVP and DVP under AO and non-AO conditions. Yellow arrows mark the analyzed structures. Magenta dashed lines indicate the half-maximum level for FWHM measurement. (B) Lateral and axial FWHM measurements of microglial processes in AO and non-AO images, showing consistently smaller process widths with AO correction. (C) The dynamic of MCs in three retinal layers at different time points from 0 min to 15 min or 30 min.

Figure S5:

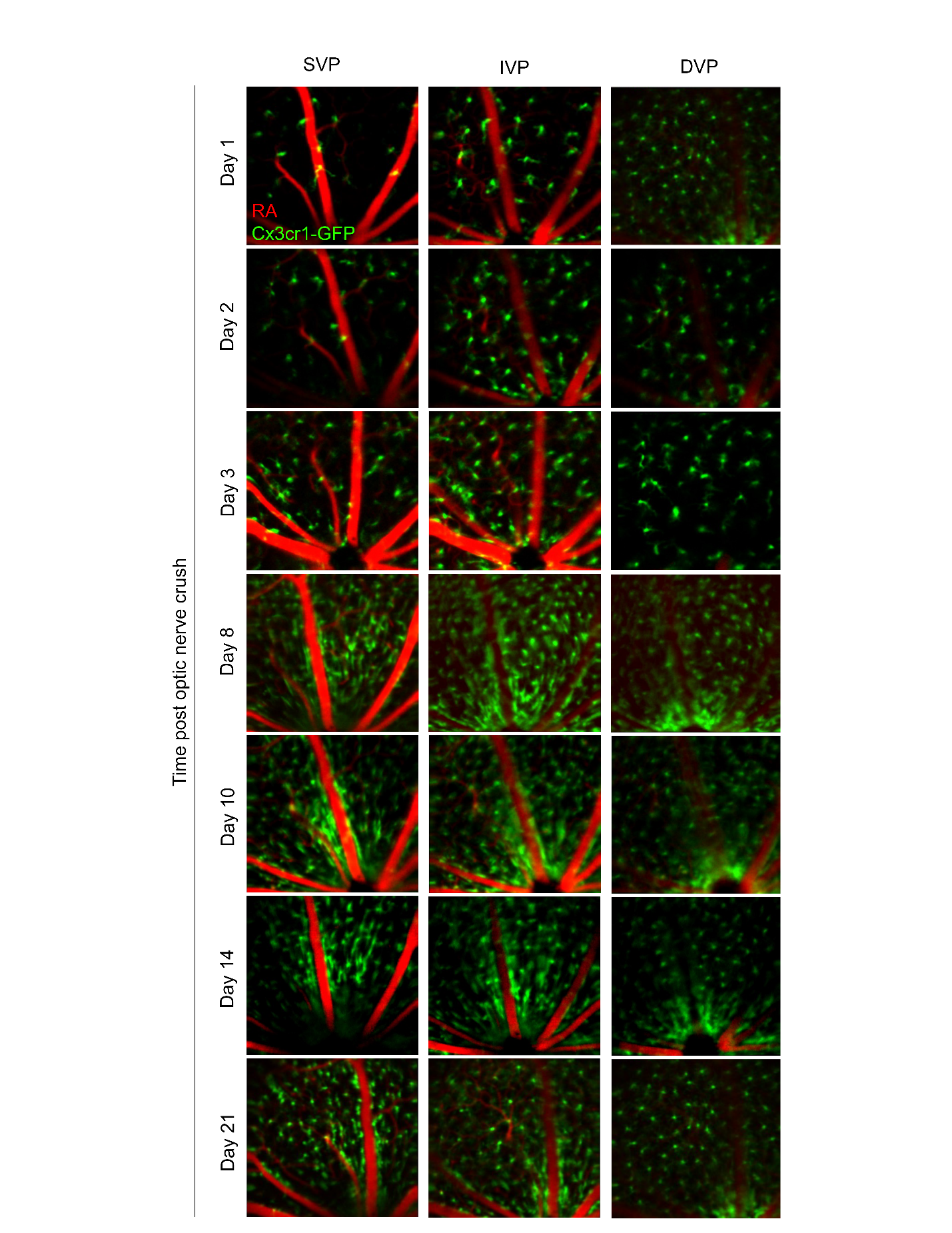

**Figure S5. The in-vivo imaging of MCs morphology in the three vascular layers on the day 1, day2, day 3, day 8, day 10, day14, and day 21 after ONC.**

Figure S6:

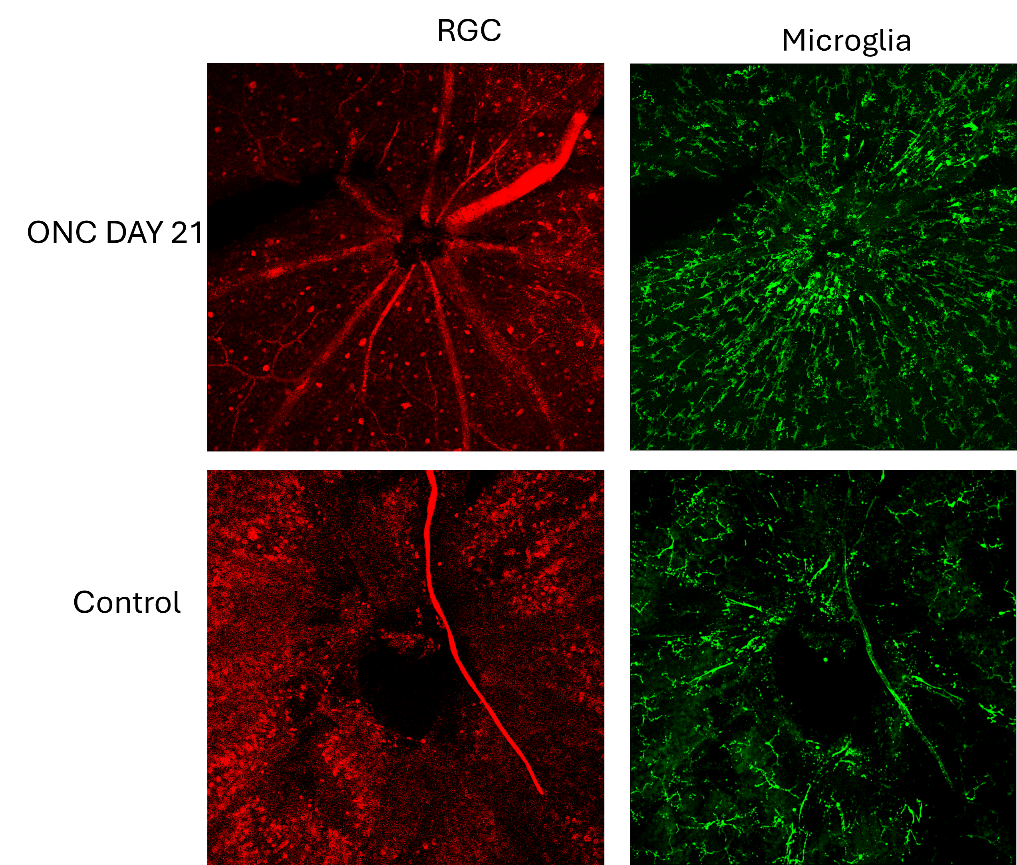

**Figure S6. The difference of RGC and microglia between ONC and control mice at SVP layers.**

Figure S7:

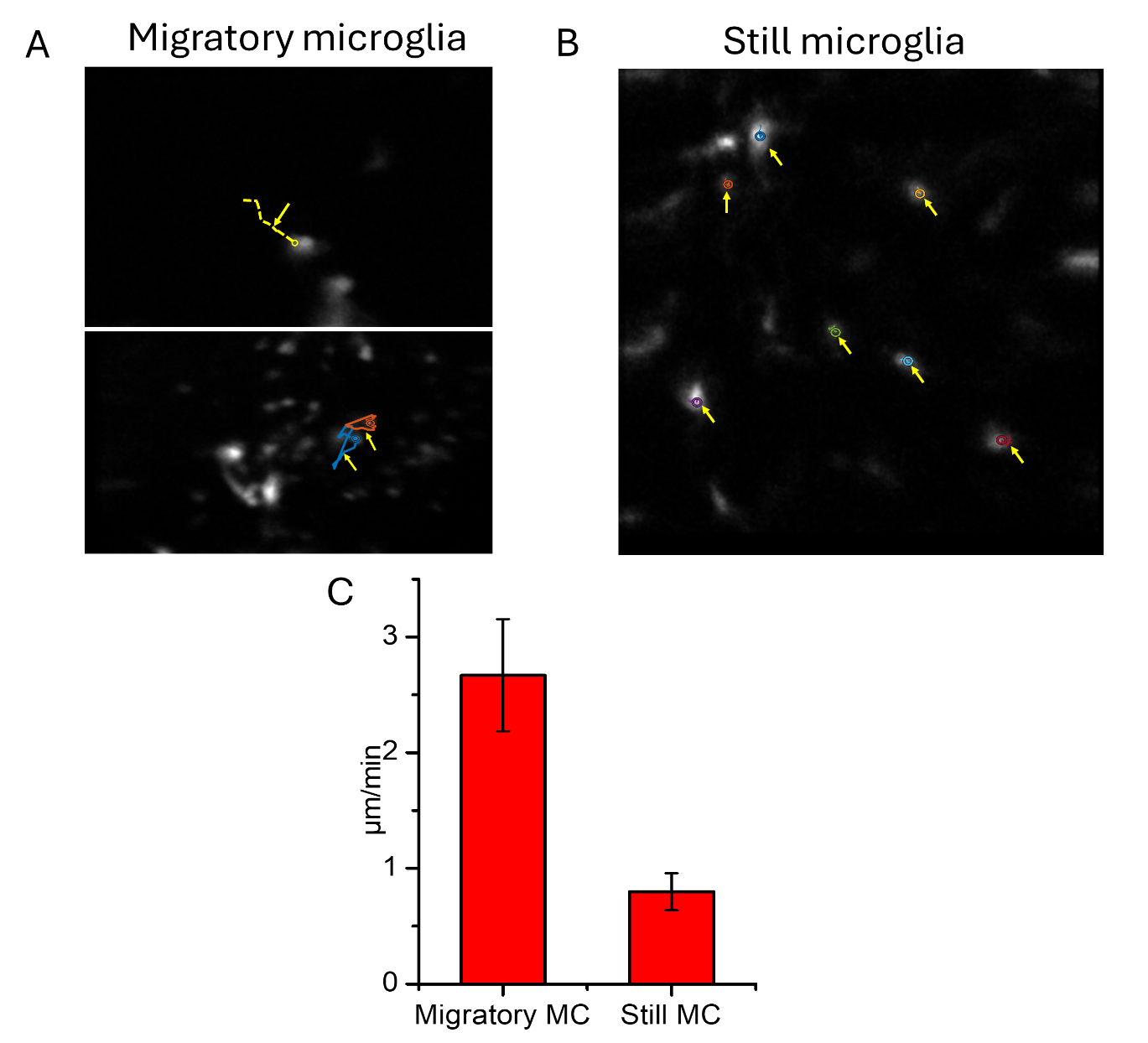

**Figure S7. The dynamic imaging of microglia cells near tRGCs after RGC transplantation.** (A) and (B) The dynamics of migratory and still microglia after RGC transplantation. (C) The corresponding migration speed of migratory and still microglia.

**Materials and Methods**

Figure S9:

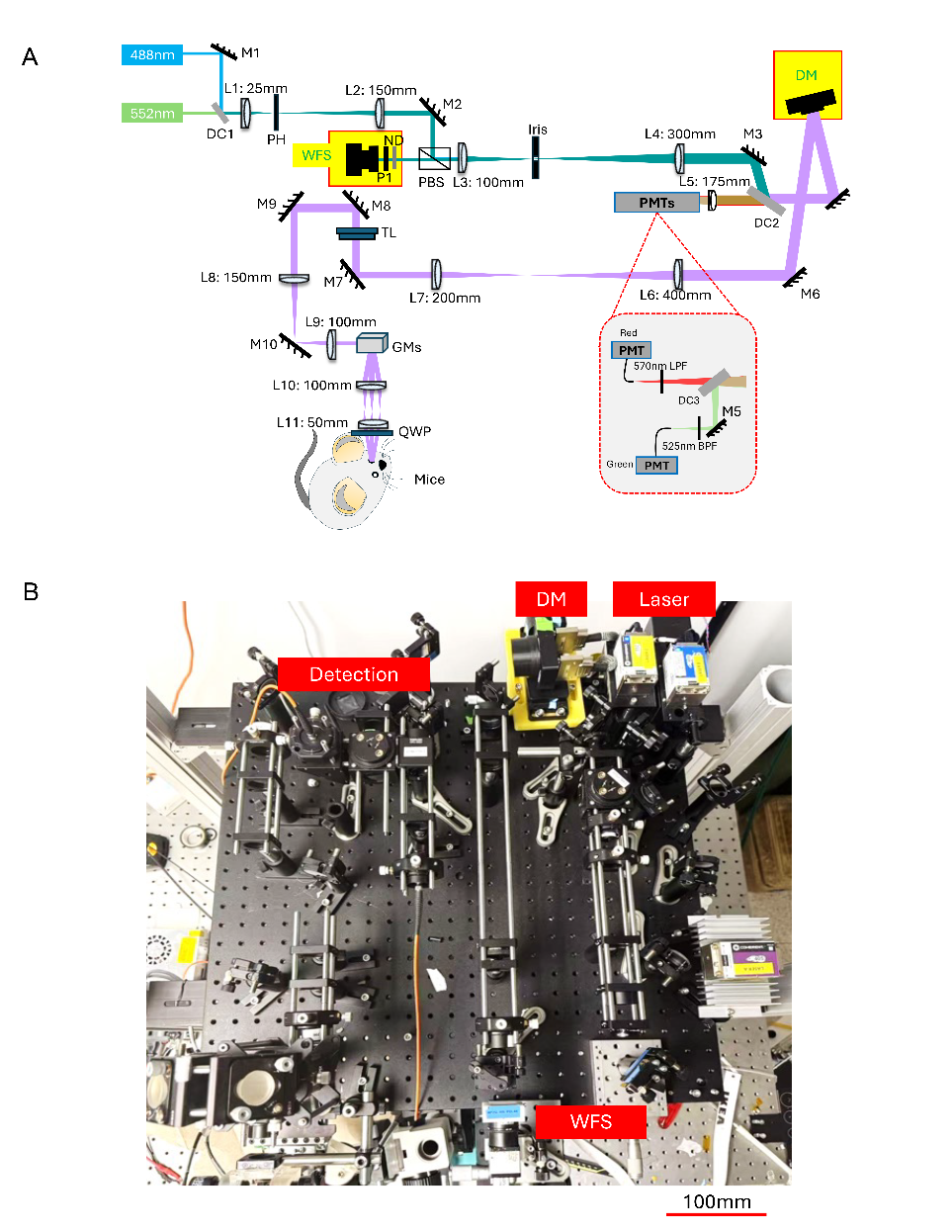

**Figure S9. (A) System schematic diagram of DC-AOSLO and (B) system layout install on the 600 mm by 600mm breadboard.** L1-L11: Achromatic Doublets Anti-Reflection Coated Lens; M1-M9: Protected Silver Mirror; DC1-DC4: Dichroic mirrors; TL: Tunable Lens; GM: Galvo Scanning mirrors; BPF: Band-pass filter; LPF: Long-pass filter; PMT: Photomultiplier tubes; PBS: Polarized Beam Splitter; DM: Deformable mirror; WFS: Wavefront Sensor; QWP: Quarter-wave plate.

Figure S10:

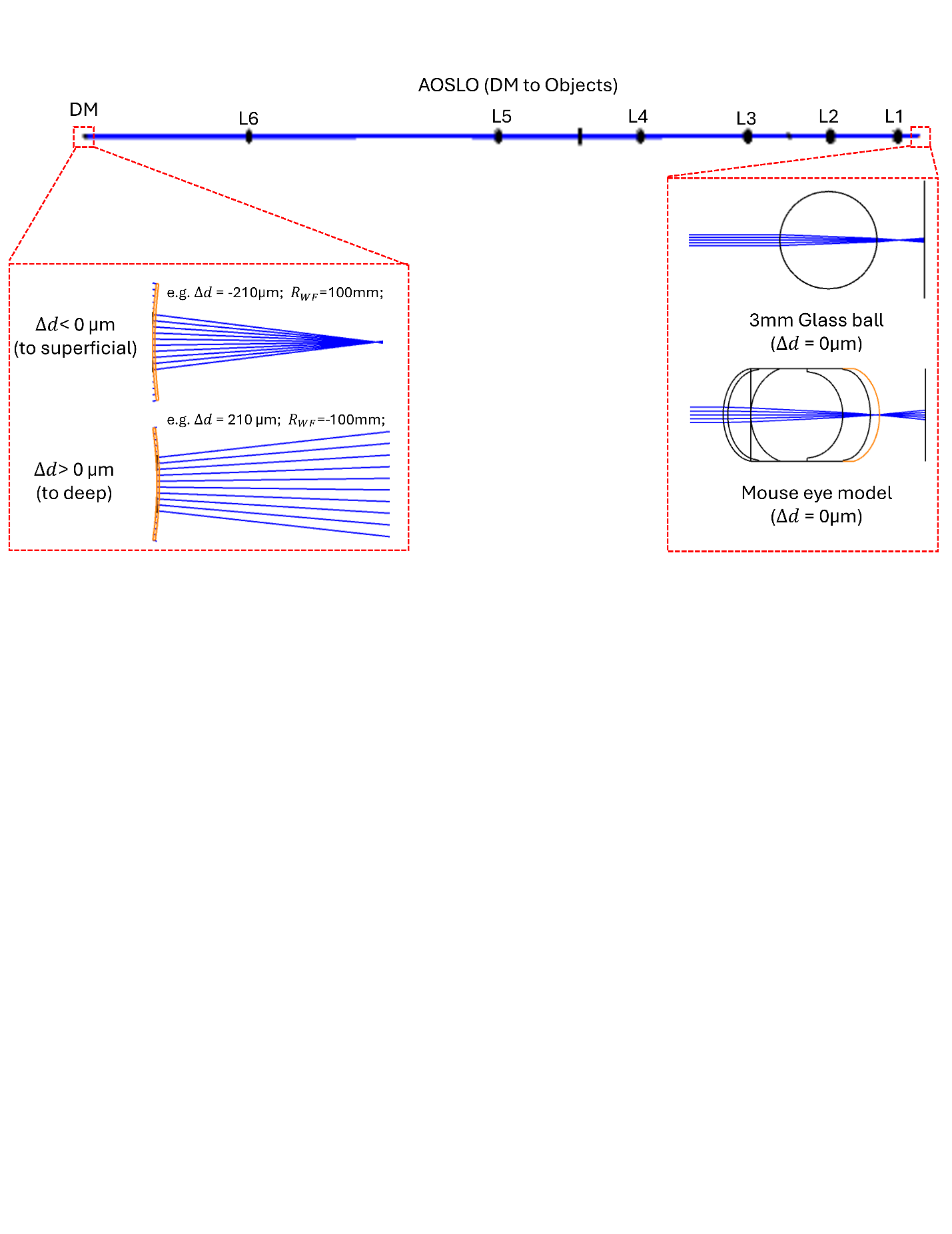

**Figure S10. Simulation of the AOSLO from imaging objects to DM for based on glass ball and mouse eye model.**

Figure S11:

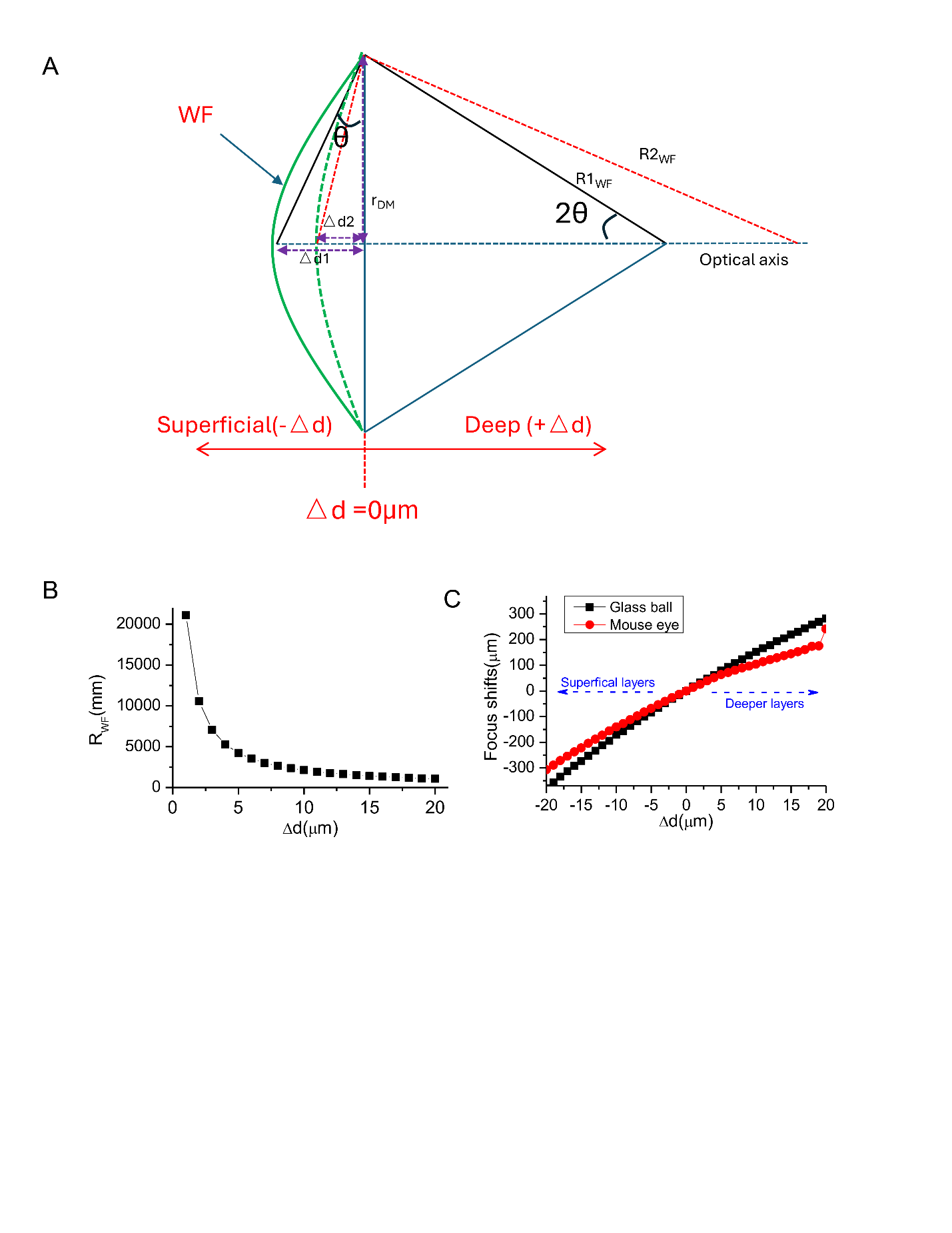

**Figure S11. The relationship between radius** $\boldsymbol{R}_{\boldsymbol{WF}}$ **of DM surface arc and DM surface shift** $\boldsymbol{\Delta d}$ **for the Zemax simulation.** (A) the math model of radius $R_{WF}$ of DM surface arc and DM surface shift $\Delta d$. (B) Radius $R_{WF}$ changes with different $\Delta d$. (C) the location changes of the image plane with different $\Delta d$.
